# Evaluating the performance of malaria genomics for inferring changes in transmission intensity using transmission modelling

**DOI:** 10.1101/793554

**Authors:** Oliver J. Watson, Lucy C. Okell, Joel Hellewell, Hannah C. Slater, H. Juliette T. Unwin, Irene Omedo, Philip Bejon, Robert W. Snow, Abdisalan M. Noor, Kirk Rockett, Christina Hubbart, Joaniter I. Nankabirwa, Bryan Greenhouse, Hsiao-Han Chang, Azra C. Ghani, Robert Verity

## Abstract

Advances in genetic sequencing and accompanying methodological approaches have resulted in pathogen genetics being used in the control of infectious diseases. To utilise these methodologies for malaria we first need to extend the methods to capture the complex interactions between parasites, human and vector hosts, and environment. Here we develop an individual-based transmission model to simulate malaria parasite genetics parameterised using estimated relationships between complexity of infection and age from 5 regions in Uganda and Kenya. We predict that cotransmission and superinfection contribute equally to within-host parasite genetic diversity at 11.5% PCR prevalence, above which superinfections dominate. Finally, we characterise the predictive power of six metrics of parasite genetics for detecting changes in transmission intensity, before grouping them in an ensemble statistical model. The best performing model successfully predicted malaria prevalence with mean absolute error of 0.055, suggesting genetic tools could be used for monitoring the impact of malaria interventions.

---

Molecular tools are increasingly being used to understand the transmission histories and phylogenies of infectious pathogens^1^. Using phylodynamic methods it is now possible to estimate the historic prevalence of infection directly from molecular data, even in organisms with relatively complex lifecycles^2^. However, these tools typically rely on pathogens having an elevated mutation rate and not undergoing sexual recombination, which allows for the application of coalescent theory^3^. Consequently, these techniques are yet to be adapted for the study of *P. falciparum* malaria, which is known to undergo frequent sexual recombination. In addition, malaria transmission between both the human and the mosquito hosts involves a series of population bottlenecks^4,5^, which combined with the brief sexual stage involving a single two-step meiotic division^6^, have marked effects on the population genetics of *P. falciparum*^7,8^. This is extenuated by evidence of cotransmission of multiple clonally related parasites^9^, which combined with host mediated immune^10,11^ and density-dependent regulation of superinfection^12,13^ result in a complicated network of processes driving the genetic diversity of the parasite population within an individual host.

Despite this substantial complexity, an increasingly nuanced understanding of the processes shaping parasite genetic diversity is appearing, with multiple genetic metrics proving promising for inferring transmission intensity^14,15^. For example, measures of the multiplicity of *P. falciparum* infections have been shown to be useful for identifying hotspots of malaria transmission^16,17^. The spatial connectivity of parasite populations has also been shown to be well predicted by pairwise measures of identity-by-descent^18,19^. More recently, it has been shown that malaria genotyping could be used to enhance epidemiological surveillance^20^, however, two main challenges have been identified before molecular tools could be used in an operational context. The first is that our understanding of the relationship between transmission intensity and within-host parasite genetic diversity is incomplete. Combined models of both population genetics and malaria epidemiology would allow us to develop a more detailed view of both processes, yet these two approaches are largely explored separately. Recent efforts have been made to incorporate both modelling scales within one framework^21^, with the concomitant modelling of resistance evolution both within and between hosts yielding important insights into the evolution of drug resistance^22^. However, the realism of either the transmission process or the genetic evolutionary process has been limited in these models, with the representation of recombination and the parasite lifecycle within the mosquito often simplified. This makes the generalisability of using molecular tools for surveillance difficult. More realistic models are subsequently needed that capture both processes. These models could answer previous hypotheses^23,24^ about how transmission intensity alters the rate at which superinfection events and cotransmission of genetically related parasites shape the parasite genetic diversity observed within humans. The second challenge is to understand in what situations molecular tools will offer advantages over traditional surveillance. In addition, power calculations need to be carried out to understand how many samples are required for reliable inference and what types of genomic data are most informative.

Here we use mathematical transmission modelling to address these challenges. We extend a previously published malaria transmission model^25^, which now allows parasite populations to be followed explicitly through the parasite’s obligate sexual life cycle by the inclusion of individually modelled mosquitoes. The new model is fitted to parasite single nucleotide polymorphism (SNP) genotype data to capture the observed relationship between an individual’s age and their complexity of infection (COI), defined as the total number of genetically distinct parasite strains in an individual. Using the fitted model, we characterise how six measures of parasite genetic diversity respond to changes in transmission intensity. We continue by conducting a power analysis, assessing the ability of each metric to detect changes in transmission intensity as a function of the number of available samples. We conclude by building an ensemble statistical model, which demonstrates how routinely collected clinical genotype samples could be used for accurate prediction of malaria prevalence using as few as 200 SNP genotyped samples.

## Results

### Complexity of Infection Data

First, we used *THE REAL McCOIL*^26^ to estimate the COI from SNP genotyped samples collected previously from individuals with evidence of asexual parasitemia by microscopy from regions in Kenya and Uganda. (Figure 1) These two datasets were selected as they recorded both the age of the sampled individuals and SNP intensities at sufficiently large number of loci, enabling the relationship between COI and age to be estimated. After excluding SNP loci with more than 20% missing data and subsequently removing samples with more than 25% missing SNP data from further analysis, the COI was estimated for 2419 samples from 95 primary schools in Western Kenya (1363 from Nyanza province and 1056 from Western province) and 584 samples from representative cross-sectional household surveys in three sub-counties in Uganda (462 from Nagongera in Tororo District, 74 from Kihihi in Kanungu District, and 48 Walukuba in Jinja District). Distribution of COI varied between each region, ranging between 1 – 21 and broadly peaking in children aged six years old before decreasing with increasing age of the individual sampled.

**Figure 1:**
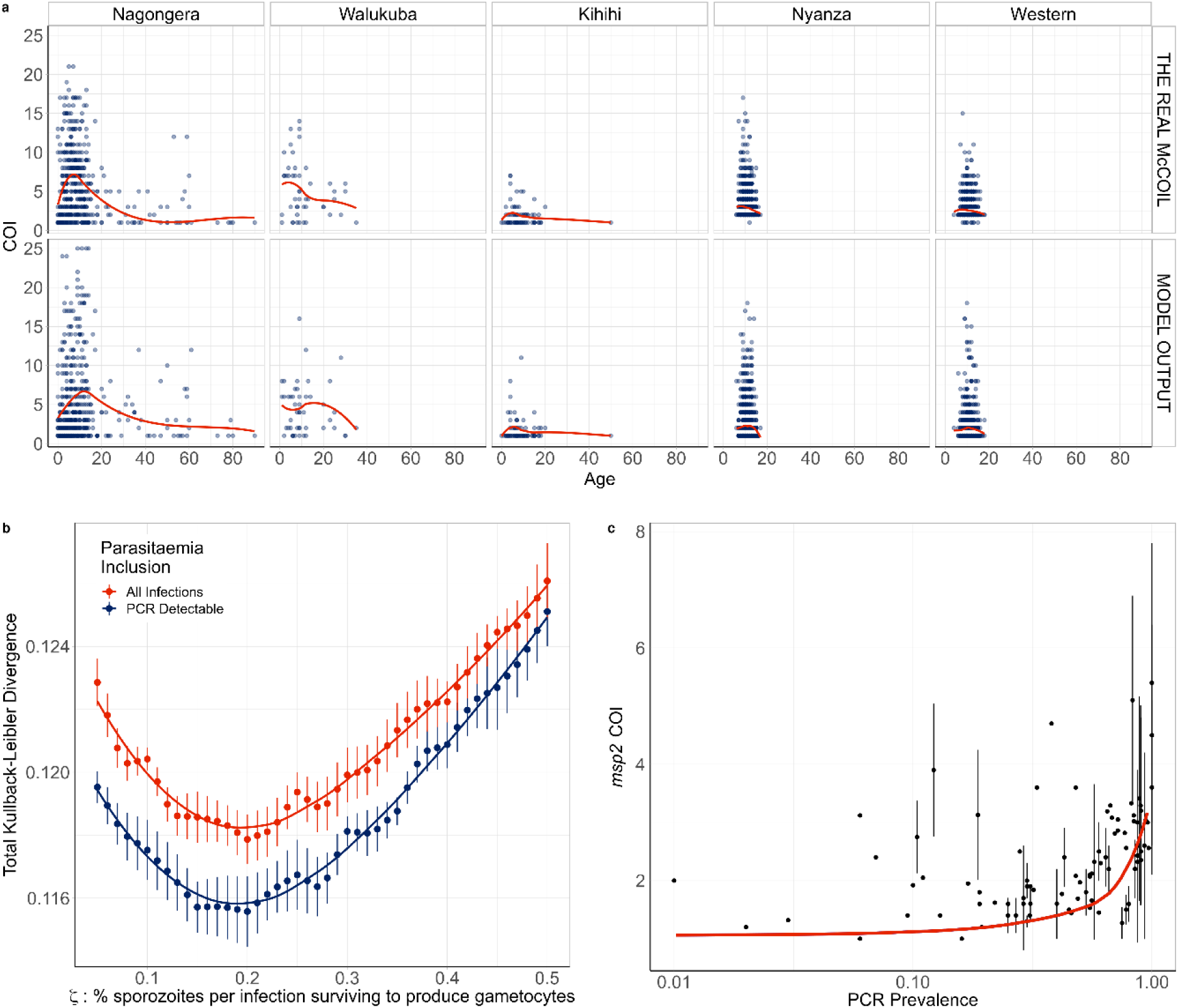
Modelled estimates of the relationship between complexity of infection against age. **a)** One realisation of the model predicted relationship between complexity of infection (COI) and age compared to the observed relationship estimated using *THE REAL McCOIL*. Each point represents an individual, with a local regression fit plotted in red. The relationship shown represents the selected best model fit, which estimates that 20% of sporozoites successfully progress to blood-stage infection in an individual with no immunity. In **b)** the results of the model fit are shown, with each point representing the mean Kullback-Leibler divergence and the whiskers representing the 95% confidence interval. Results of model fitting are shown for the assumption that all infections are detected (red) or only those that are PCR-detectable (blue). In **c)** the model predicted relationship between COI measured by *msp2* genotyping and PCR prevalence is shown in red, with the point-ranges showing observed values of COI by *msp2* genotyping from the literature review.

### Fitted Model

We developed an extended version of a previously published individual-based model of malaria transmission^25^. Briefly, the model was extended to include individual mosquitoes, enabling parasite populations and their genotypes to be tracked throughout the full lifecycle, enabling the potential formation of multiple oocysts from an infectious event and multiple genetically distinct sporozoites to be onwardly transmitted. Male and female gametocytes are sampled from the infecting human with the probability proportional to relative densities of each genotype. The resultant oocyst is able to produce up to four new parasite genotypes resulting from a two-step meiotic division. The extensions require use to define the proportion of sporozoites from an infectious bite that survive to found a blood stage infection, which we define as *ζ*. This process will ultimately affect the level of new parasite genetic diversity introduced and consequently we parameterised our developed model (see Materials and Methods and Supplementary methods) through fitting to the earlier estimated relationships between COI and age in the five regions across Uganda and Kenya (Figure 1a). We estimate that 20% of sporozoites onwardly transmitted within an infectious bite successfully progress to a blood-stage infection and produce gametocytes that may contribute to future mosquito infections. The model captures the observed peak in COI observed at age 7-8 (Figure 1a); however, the comparatively fewer samples at higher ages make it difficult to confirm that this is the true peak in COI. Additionally, this observed peak in COI also likely reflects the limits of detection, with more accurate model predictions occurring under the assumption that parasite strains that would not be detected by PCR do not contribute to the estimated COI (Figure 1b). Model fitting also showed that sensitivity of the model fit to the percentage of sporozoites that survive is negligible between values of 15-20%, with the confidence intervals for the most likely parameter value of *ζ* overlapping intervals for values of *ζ* ranging from 0.1 to 0.29.

To further assess the fitted model, we wanted to incorporate estimates of COI based on *msp2* genotyping, which is more commonly measured, however, it does underestimate COI in individuals with high COI, with COIs > 7 difficult to resolve. We updated a previous literature review^16^ of paired estimates of *msp2* COI and parasite prevalence by PCR, which yielded 91 paired measures of *msp2* COI and PCR prevalence. The fitted model predicts an increase in *msp2* COI with increasing malaria prevalence in agreement with the data collected within our literature search (Figure 1c). However, there are notably larger uncertainties in the recorded *msp2* COI at higher prevalence ranges in the studies found.

### Contribution of cotransmission events to within-host parasite diversity

Using the fitted model, we explored the relationship between the proportion of within-host parasite strains that are highly-related, which we define as being more than 50% IBD with other parasites and thus indicative of cotransmission events, and transmission intensity. The model-predicted proportion of within-host parasite diversity that is due to cotransmission events was shown to increase at lower transmission intensities (Figure 2a). We predict that at PCR prevalence less than 11.5%, more than 50% of strains within polygenomically infected individuals of all ages result from cotransmission events, rather than superinfection. This is based on the assumption that highly related parasites have originated from a recent common ancestor, and as such reflects the proportion of within-host genetic diversity that is due to cotransmission events rather than superinfection. We also predict this relationship is dependent on the age of individuals sampled, with parasites within younger individuals more likely to be more highly related. This reflects the increased chance that younger individuals will be treated after an initial infection due to their lower acquired immunity increasing the probability of developing clinical symptoms from an infection. Subsequently, younger individuals will be less able to accrue parasites from superinfection events, which increases the likelihood that any polyclonal individuals are the result of a cotransmission event. In Figure 2b, the model-predicted relationship between mean IBD in mixed infections and the fraction of mixed infections is shown, and is well described by an exponential trend line fit to this data. The model-predicted relationship is comparable to estimates of IBD from whole genome sequence data collected from sites across Africa and Asia as part of the Pf3k study^27^. However, the model predicts significantly lower mean IBD in settings with a high fraction of mixed infections compared to the estimates based on the whole genome sequencing data, with samples from sites in Ghana, Malawi, Mali and the Democratic Republic of the Congo exhibiting higher mean IBD than predicted by the model.

**Figure 2:**
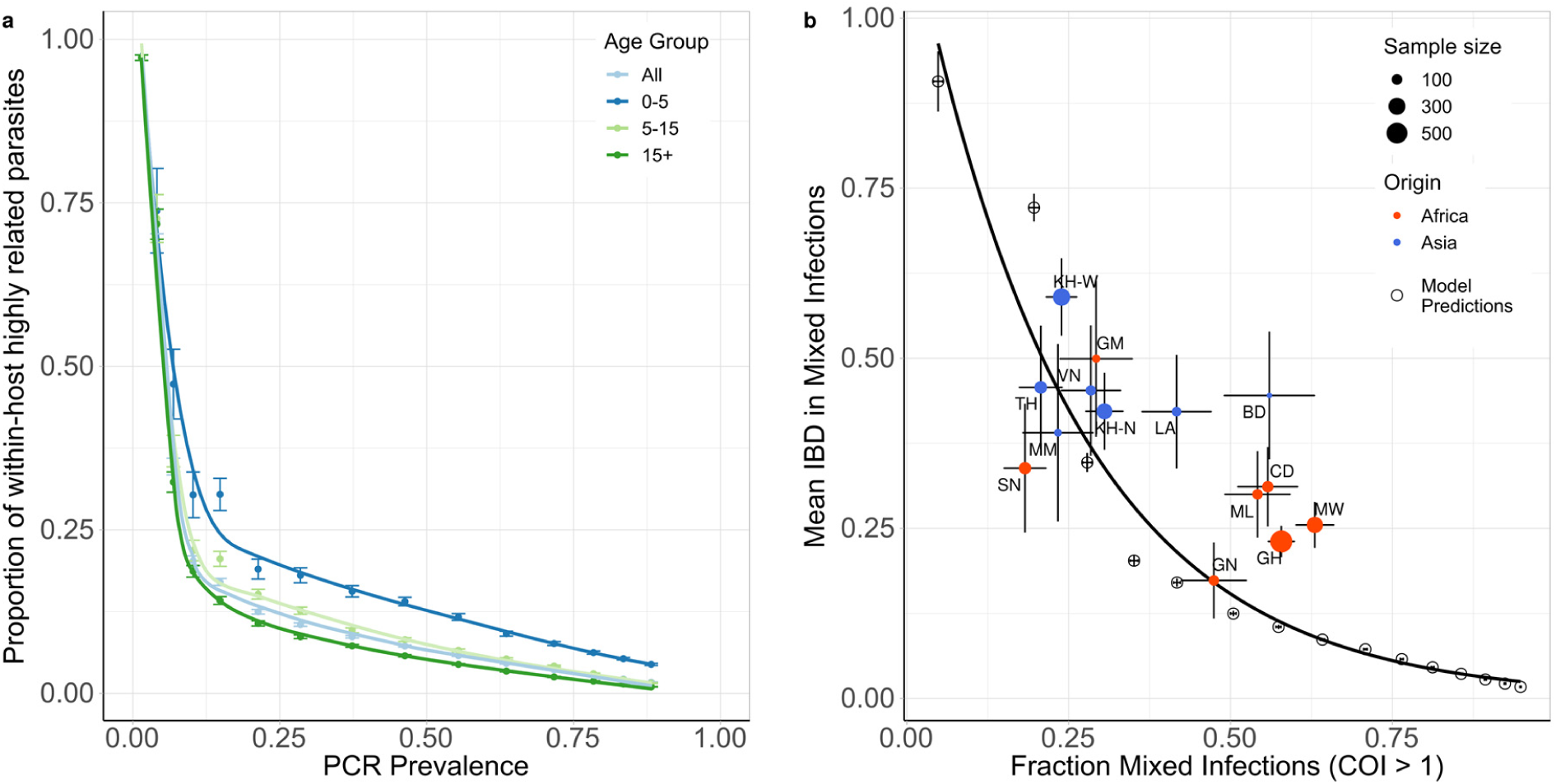
Contribution of superinfection and cotransmission to within host parasite relatedness. In **a)** the model predicted relationship between the mean within host proportion of highly identical parasite strains (>50% of loci comparisons are identical by descent (IBD)) against PCR prevalence. The relationship is shown for all ages and for three age groups: 0-5 years, 5-15 years and 15+ years, with error bars showing ±1 standard error of the mean. In **b)** the mean IBD in mixed infections (COI > 1) is shown against the proportion of mixed infections. Results from model simulations are shown with empty circles with an exponential regression shown with the black curve. The model estimates are compared to estimates of IBD from whole genome sequence data collected in sites across Africa and Asia, which were estimated previously in Zhu et al^27^. Populations are coloured by continent, with size reflecting sample size and error bars showing ±1 standard error of the mean. Abbreviations: SN-Senegal, GM-The Gambia, NG-Nigeria, GN-Guinea, CD-The Democratic Republic of Congo, ML-Mali, GH-Ghana, MW-Malawi, MM-Myanmar, TH-Thailand, VN-Vietnam, KH-Cambodia, LA-Laos, BD-Bangladesh.

### The impact of intervention strategies on parasite genetic diversity

Using our parameterised model, we first modelled how a reduction in transmission would affect four genetic metrics as the prevalence of malaria declined due to the scale up of interventions (Figure 3). The genetic metrics explored were: 1) the population mean complexity of infection (COI), 2) the % of samples that are polygenomic (COI > 1), 3) the % of unique parasite 24-SNP barcodes and 4) the coefficient of uniqueness (COU) (Figure 3). COU is a new measure of genetic relatedness within samples and is equal to 0 when all barcodes within a sample are identical, and is equal to 1 when all barcodes within a sample are unique (a multi-locus analogue of homozygosity).

**Figure 3:**
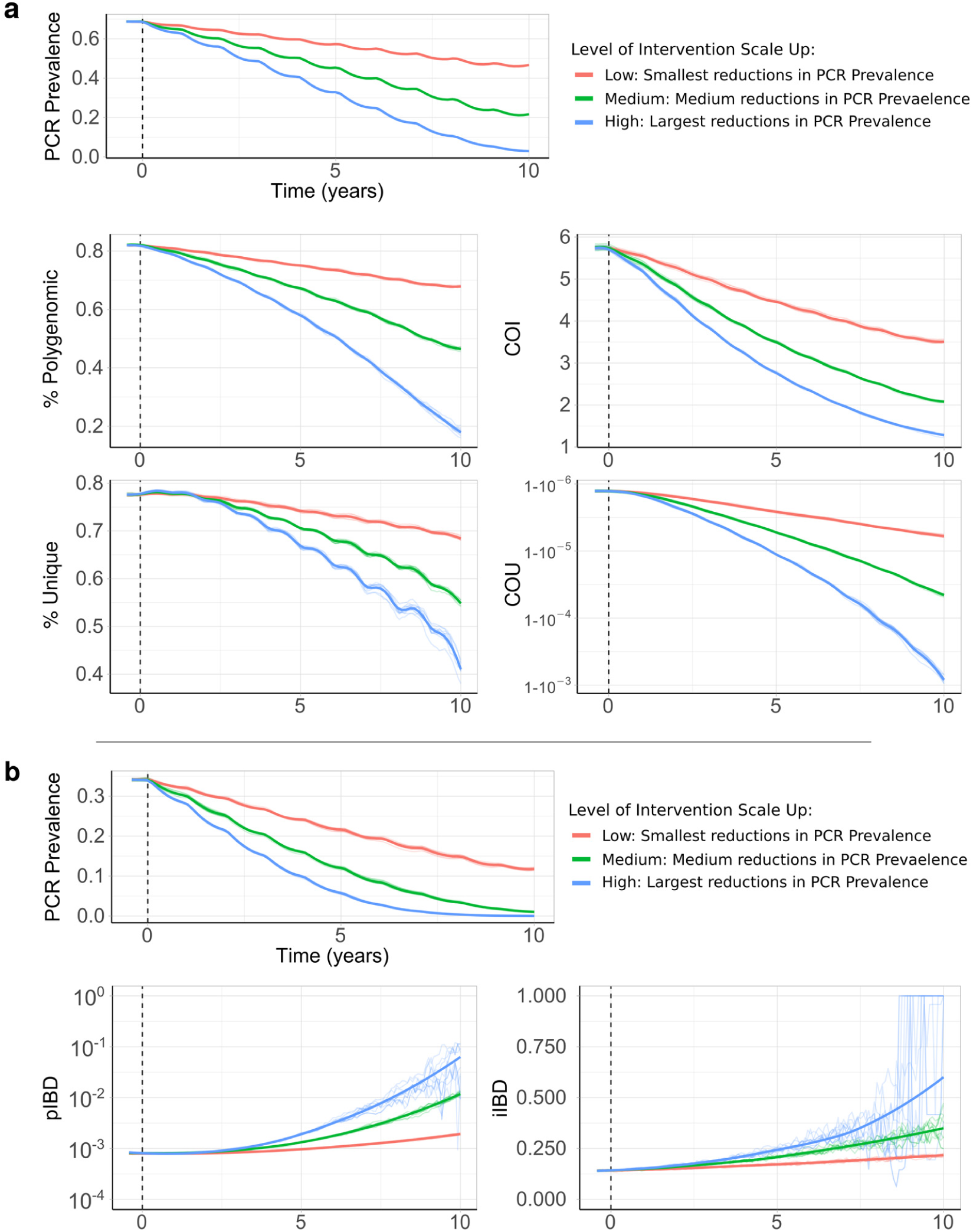
Impact of changes in transmission intensity upon genetic metrics of transmission intensity. In **a)** the top plot shows the change in PCR prevalence after the introduction of 3 different levels of intervention scale up, with both the 10 individual stochastic realisations and the mean local regression smoothed relationship shown. The following four plots show the population mean percentage of the population that are polygenomically infected, the complexity of infection (COI), the percentage of samples that are genotypically unique (% unique) and the coefficient of uniqueness (COU) for the prevalence declines seen in the first row. In **b)** the top plot shows the change in PCR prevalence, which starts at a lower starting prevalence of 35% compared to 70% in **a)**. The following row shows the within-host identity-by-descent (iIBD) mean across the 24 identity loci considered, and the population mean pairwise measure of IBD (pIBD). In all plots the vertical dashed black line shows the time from which the scale up of interventions starts (Time = 0 years).

The model was initiated at 70% PCR prevalence with no interventions in place. Three levels of intervention scale-up were simulated, representing a low, medium and high reduction in prevalence resulting in a final PCR prevalence of ∼45%, ∼20% and ∼5% respectively after ten years. We predict that all four metrics decline proportionally with declining malaria prevalence (Figure 3a). The model predicts that the specific relationship depends on the population chosen for genetic testing (Supplementary Figure 1a). For example, COI is predicted to be higher in older age categories. The percentage of unique samples varied greatly depending on the on the sub-population sampled, reflecting difference in the absolute numbers of individuals that fall within each sub-population. Samples taken from individuals with asymptomatic infections were predicted to have the highest COI and percentage of polygenomic samples. Across the scenarios simulated, metrics based on the complexity of infection (COI and % Polygenomic) showed a higher level of correlation with changes in the prevalence of malaria than measures based on the uniqueness of samples (COU and % Unique) (Table 1). In addition, samples collected only from patients with symptomatic malaria led to metrics that were the least correlated with reductions in prevalence, resulting from the decreased number of available samples. This effect was most noticeable when assessing the percentage of unique genotypes within clinical samples, which had a correlation coefficient of 0.24 with PCR prevalence (Table 1).

**Table 1:**
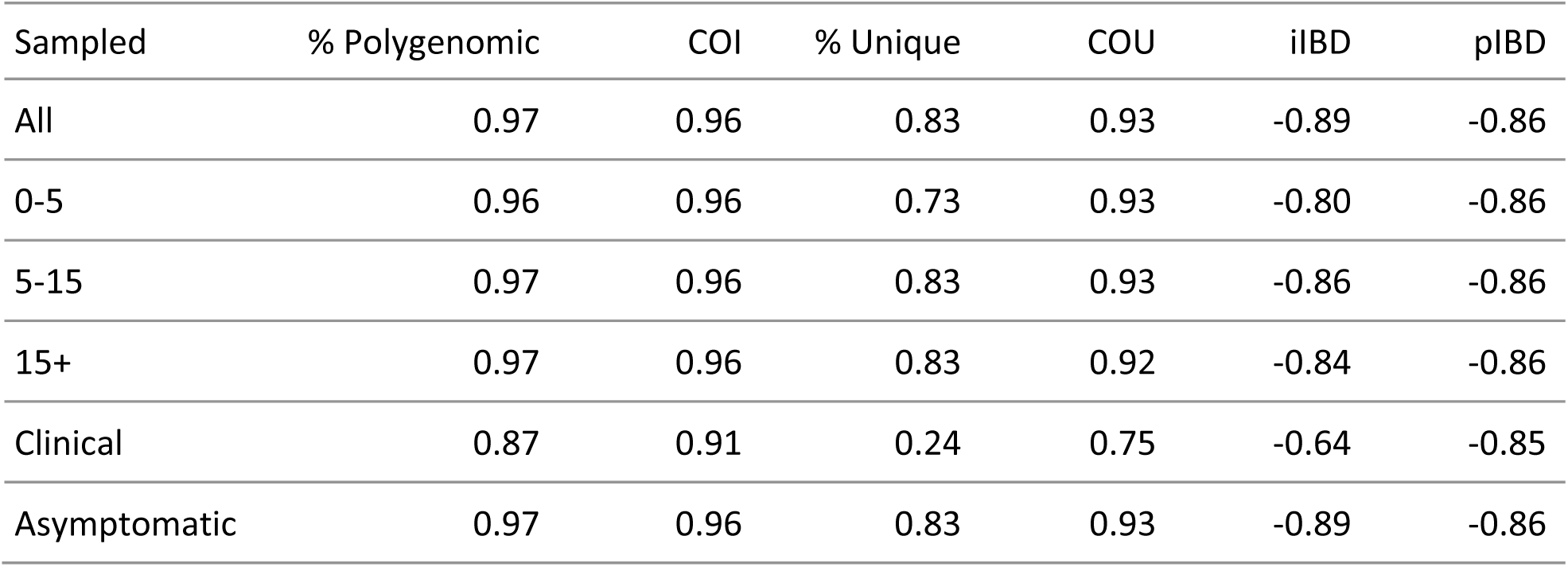
Kendall rank correlation coefficients between genetic diversity metrics and parasite prevalence. Coefficients are bound between −1 and 1, with 1 indicating perfect ranked positive correlation and −1 indicating perfect ranked negative correlation.

We also assessed measures of parasite genetic diversity based on comparisons of the number of loci that are identical-by-descent (IBD), which included the within-host pairwise mean proportion of loci that are IBD (iIBD) and the population pairwise mean proportion of loci that are IBD (pIBD). We predict that both metrics increase in response to declines in prevalence, however, we predict that pIBD only increases substantially at PCR prevalences less than 15% (Figure 3b). Consequently, metrics based on IBD were explored at a lower starting prevalence of 35% PCR prevalence before the scale up of interventions. The shape of the increase in iIBD was predicted to be dependent on the population sampled (Supplementary Figure 1a), with iIBD increasing quicker in symptomatic individuals. iIBD, however, becomes less informative as transmission intensity declines, with individuals less likely to be infected with multiple strains due to the lower rates of superinfection.

### Power Analysis

To evaluate the performance of each metric for detecting annual changes in the prevalence of malaria, we calculated the statistical power for each metric at different sample sizes, focussing on samples collected from children aged between 5-15 years old. We estimate that after 5 years of intervention scale up, corresponding to an absolute decrease in malaria prevalence by PCR of 20%, no more than 350 samples are required for each metric explored (except for iIBD) to detect the change in transmission intensity 80% of the time (Figure 4). The predictive power, however, declined across all metrics when the effect size, i.e. the decrease in prevalence, decreased. With 600 samples, each metric had less than 40% power to detect the decrease in prevalence after 1 year. The performance of each metric was additionally dependent on the starting prevalence, with metrics based on the uniqueness of samples (COU and % Unique) predicted to be more powerful at lower starting prevalences compared to higher prevalences (Figure 4b). Metrics based on measures of IBD were overall less powerful, with the predictive power of iIBD being less than 80% across all years and sample sizes (Figure 4c). pIBD only exhibited a predictive power greater than 80% when detecting the largest change in prevalence between 22.5% and 8%, requiring over 225 samples.

**Figure 4:**
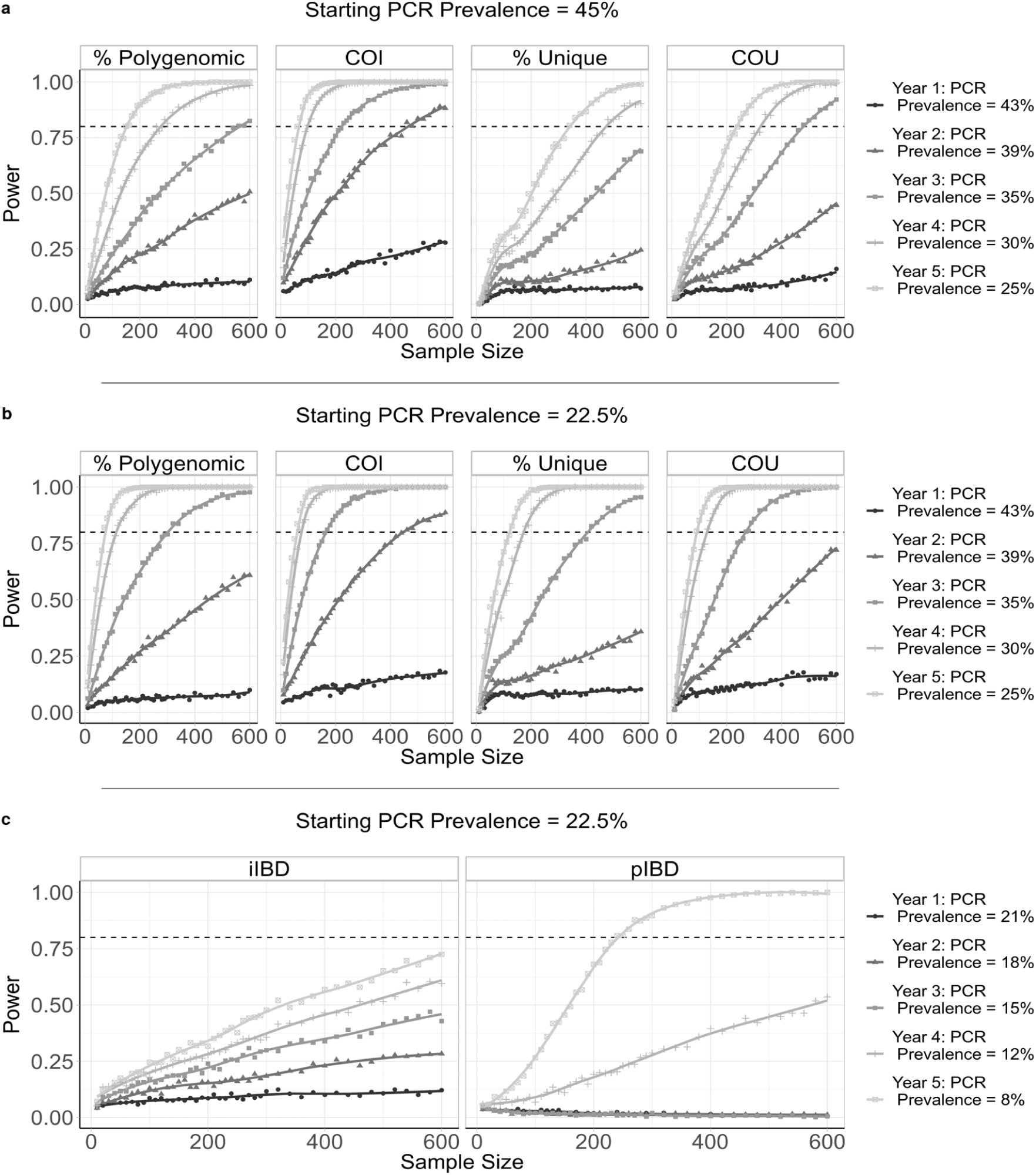
Predictive power of six metrics of parasite genetic diversity with respect to sample size. The distribution of sample means of six metrics of parasite genetic diversity were compared for five years following the initiation of the scale up of intervention coverage. For each sample size, the power is defined as the proportion of 100 subsamples comparing year 0 and years 1-5 for which a significant difference in the mean was observed, estimated using one-tailed Monte Carlo p-values generated by 100 permutations of the years samples were collected in. In **a)** the metrics assessed are the percentage of samples that are polygenomic, the complexity of infection (COI), the percentage of barcodes within samples that are unique, and the coefficient of uniqueness (COU). The power of each metric was compared across five years in which a 20% absolute decrease in parasite prevalence from 45% was observed. The same information is shown in **b)**, but for a 14.5% absolute decrease in prevalence from 22.5% over 5 years. In **c)** the metrics considered are the mean withinhost identity-by-descent (iIBD) and the population mean pairwise measure of IBD (pIBD). In each plot 80% power is shown with the horizontal dashed line.

The power of COU, % Unique and pIBD were noticeably worse when it was assumed that samples from polygenomically infected individuals could not be phased (Supplementary Figure 2). Under this assumption we assume that we are unable to observe the genotype of each strain and consequently only the major haplotype within an individual is available, i.e. calling the most abundant allele at each locus of the barcode, which negates our ability to measure an individual’s iIBD. Across the full range of malaria prevalence simulated, measures of COI and COU were consistently predicted to be the most powerful, with % unique samples and IBD metrics demonstrating increased power to detect changes in transmission in areas with lower baseline transmission intensities where we predict the genetic variation to be lower.

### Statistical model for predicting transmission intensity

In order to translate the information we have characterised into an effective tool for assisting surveillance programs, a statistical model was created to predict malaria prevalence using genomic metrics derived from parasite SNP genotyping (see Materials and Methods). Due to the difficulty in phasing high complexity infections, we assumed that all collected samples were unphased and as such we did not focus on metrics based on IBD when building our data set for training our statistical model.

The fitted ensemble model performed well on out-of-sample simulation datasets, and was able to identify the underlying model behaviour used to generate the training dataset (Figure 5a). The best performing model provided accurate predictions of malaria prevalence when tested on SNP genotype data from the five administrative regions, with an observed mean absolute error equal to 0.055 for these five locations. The performance of the model was enhanced when sample metadata was available (Figure 5b), with the ensemble model trained and tested using data with no age or clinical status information consistently performing worse. Similar patterns were also observed when assessing the performance of each of the level 1 models in the ensemble model (Supplementary Table 1). As in the power analysis, across the range of malaria transmission intensities assessed, measures of COI and COU were observed to be the most informative metrics (Supplementary Figure 3). Model predictors based on the age and clinical status of individuals sampled contributed 28% towards the total model importance.

**Figure 5:**
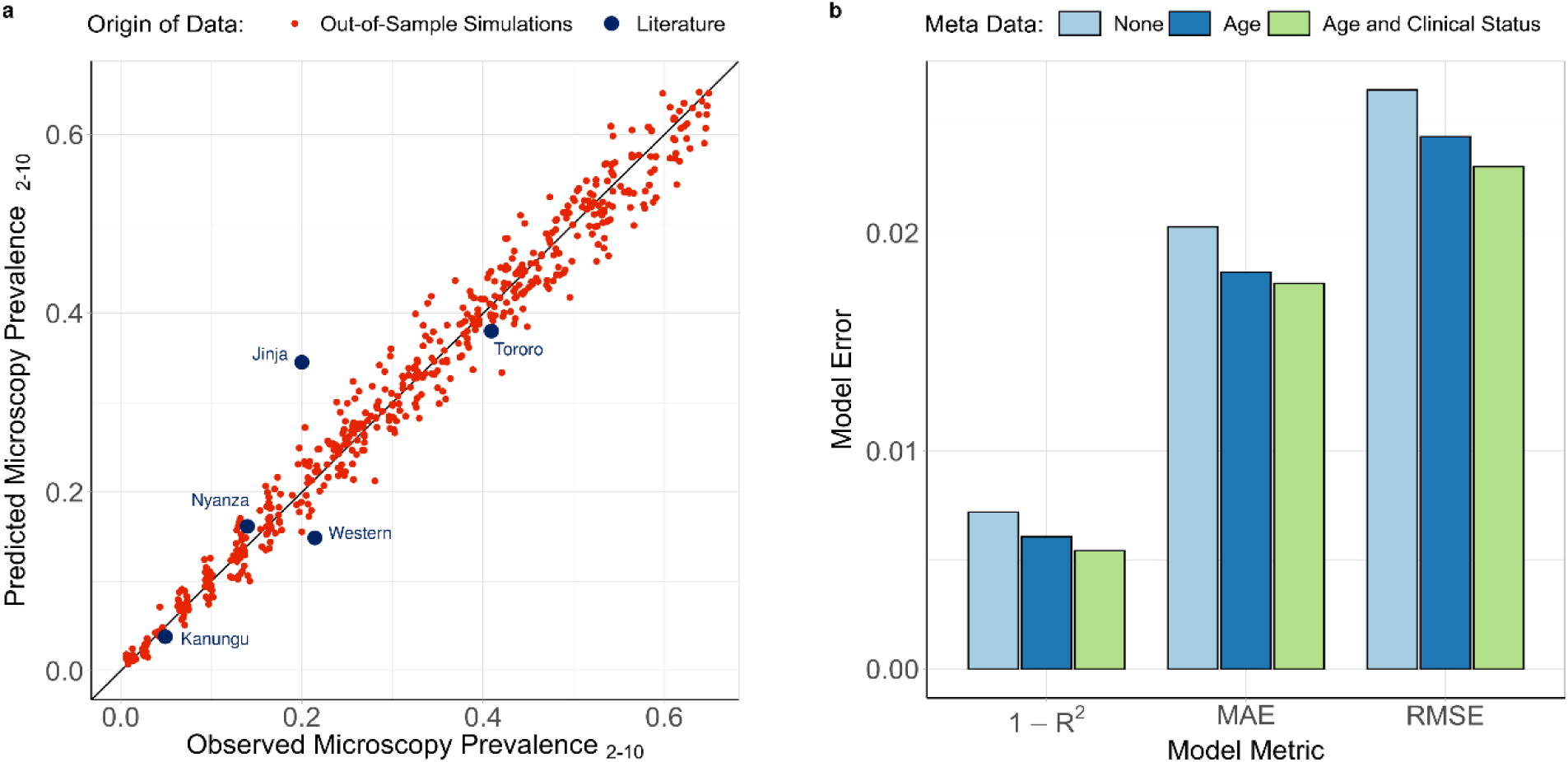
Ensemble statistical model predicted malaria prevalence vs observed malaria prevalence. In **a)** the performance of the trained ensemble statistical model is shown, with the model predicted prevalence in red showing the predictions for the out-of-sample test dataset composed of model simulations held back from model fitting. The blue points show the predicted prevalence for the 5 administrative regions considered earlier. In **b)**, the performance of the ensemble model is shown under different assumptions about the availability of patient metadata within simulated data.

## Discussion

The substantial reduction in the cost of generating genomic datasets over the last ten years and the establishment of scientific networks committed to generating and sharing genomic data has resulted in an abundance of sequenced *Plasmodium falciparum* genomes. This effort has resulted in the identification of loci associated with emerging drug resistance mechanisms^28^ and assisted in developing putative novel drug targets^29^. Another potential use of malaria sequencing efforts is understanding how malaria genomes can be used to study transmission. Simple population genetics principles predict that in a closed population a reduction in transmission intensity will typically be accompanied by a reduction in parasite genetic diversity, resulting from reduced opportunities for outcrossing to occur within the sexual stages of the parasite’s life cycle. However, there is as yet no consensus in the use of parasite genetics for inferring transmission intensity. There is a need to understand the contribution of superinfection and cotransmission towards the within-host parasite genetic diversity, which is often highlighted within critiques of early attempts to utilise modelling approaches for transmission intensity inference^30^.

In this study we have extended a previously developed model of malaria transmission to include individual mosquitoes and discrete parasite populations. The percentage of sporozoites that are successful within an infectious bite was estimated to be 20% (95% CI 10%-29%), and was estimated by fitting our model to 3002 measures of the complexity of infection and age of individuals in 5 sites across Kenya and Uganda. The fitted model was used to initially estimate the proportion of the within-host parasite genetic diversity that is the result of cotransmission events resulting in the acquisition of highly identical parasite strains, as opposed to strains acquired through superinfection events. We predict that for malaria prevalence greater than 11.5%, the majority of genetic variation within-hosts is generated through superinfection events. To our knowledge this is the first attempt to characterise this relationship across the full transmission intensity spectrum seen within sub-Saharan Africa and represents a move towards standardising which genomic metrics should be used at different transmission ranges.

We predict that IBD within samples decays exponentially as the proportion of samples is increasingly polygenomic. This exponential relationship was similar to findings in a recent study of IBD, which used whole genome sequence data to explore this relationship^27^. However, the model predicted significantly lower IBD at higher transmission settings (settings with a higher fraction of mixed infections) than observed in the data presented in Zhu et al. There are a number of reasons for this. Firstly, the whole genome sequence data was collected from individuals of unknown age as part of a convenience sample. If the samples were collected exclusively from younger individuals, the results in Figure 2a would suggest that the mean IBD would be higher than if the samples were collected across all ages. Secondly, in the study by Zhu et al, the estimated COI across all sites was less than 2, which is significantly lower than COI estimates from the sites in Kenya and Uganda in Figure 3.3. Given that some of the African study sites in Zhu et al are in areas of high transmission intensity, it seems likely that the convenience sampling scheme used has selected for individuals with lower COIs. One explanation could be that the individuals chosen for sequencing receive treatment more regularly, which reduces the probability of parasite strains from superinfection events being present at the time of sampling. This could be due to their age, or due to their enrolment in the study that resulted in them being selected for sequencing. Ultimately, without this information it is challenging to draw strong conclusions about the validity of the model predictions in Figure 3.2b, although the broad similarity is encouraging.

Our newly defined measure of parasite diversity, the coefficient of uniqueness (COU), alongside COI were consistently powerful statistical tools for detecting changes in malaria prevalence. This is hardly surprising, as we should consider that the % unique samples and the % of polygenomic samples are simply the extreme cases of these metrics, and so we would expect them to contain less information. Additionally, the power analysis conducted was under the assumption that all samples that could be detected by PCR can be effectively phased. This is an overly ambitious assumption, and it is more correct to assess these metrics under the assumption that polygenomic samples cannot be phased (Supplementary Figure 2). However, the increase in statistical power when we are able to phase samples should highlight a need within the research field for methods to compare unphased parasite samples, with the majority of samples at higher transmission intensities predicted to have a COI greater than 1.

In the absence of being able to phase polyclonal samples, however, the observed genomic metrics were still informative within the ensemble statistical model developed to translate parasite genetic information into estimates of malaria prevalence. For example, variable importance was observed for each predictor variable (Supplementary Figure 3), however, COU and COI accounted for nearly half the variance explained. There is also a degree of compensation afforded between metrics, i.e. where one metric becomes less informative, another metric becomes more predictive. For example, at PCR PfPR less than 10%, COI and the % of samples that are polygenomic will become substantially less informative, whereas IBD measures will start being more informative. This is further demonstrated by only needing 200 samples within our statistical ensemble model to produce accurate predictions of the prevalence of malaria, with the addition of individual level metadata yielding further gains in model performance (Figure 4b). As more samples are added only modest improvements in model predictive performance are observed (Supplementary Figure 4). The importance of meta data, specifically the age of individuals, is highlighted in the findings of the model predicted COI between age groups. In Figure 3, we compared the COI between asymptomatic and symptomatic individuals, in which we predicted across all ages that asymptomatic individuals have higher COI. However, this finding does not hold when we compare the COI between symptomatic and asymptomatic individuals at different age groups and across different transmission intensities. For example, in the model fitting in lower transmission areas younger children who are symptomatic are predicted to have higher COI than asymptomatic younger children (Supplementary Figure 5). This finding is reversed, however, at higher transmission intensities reflecting the interaction between acquired clinical immunity and rates of superinfection.

This study has some important limitations. Firstly, we assumed there is only one parameter detailing the percentage of sporozoites that successfully progress to a blood-stage, which is the same for all study sites considered. This is likely a simplification, but our observation of 20% sporozoites surviving from an individual mosquito feed is comparable to Bejon et al’s observation of 25% (14 sporozoites surviving from an assumed total of 55 sporozoites resulting from five mosquito bites) of sporozoites successfully progressing to blood-stage infection^31^. It is, however, higher than estimates based on transmission efficacy studies^32^. The model fitting, however, revealed that the sensitivity to this parameter was low, with the confidence intervals for a value of *ζ* equal to 0.20 overlapping intervals for values of *ζ* ranging from 0.1 to 0.29. This is highlighted when we re-examined the model predicted relationship between *msp2* COI and prevalence with these values, which showed only slight changes to the predicted COI (Supplementary Figure 6). The fitted estimate was also based on model fits to the administrative mean prevalence as opposed to the recorded prevalence in the specific study sites. For example, the study site in Jinja District, Walukuba, was observed to have the lowest parasite prevalence of all three study sites in Uganda^33^. If we had used this prevalence value as opposed to the administrative prevalence value, the parameterised model would have failed to predict the pattern of COI in Walukuba (Supplementary Figure 7), which may suggest that this study site exhibits higher heterogeneity in the force of infection. However, the fact that the model-predicted COI closely matches the observed data when using the administrative region’s prevalence may suggest that parasite genetic metrics are more representative of the prevalence at larger spatial scales, which in turn may reflect human mobility between areas of differing transmission intensity and parasite genetic diversity. This may also be of benefit from a surveillance point of view, with 200 samples being able to give accurate measures of malaria prevalence within a large area. This could be of particular utility in areas where community surveillance is not feasible, in which samples collected from symptomatic patients attending public health facilities could provide additional information in helping to translate clinical incidence into measures of parasite prevalence.

Secondly, we did not explicitly model the scale-up of vector based interventions, instead incorporating the effects of insecticide treated nets and indoor residual spraying through their impact on the average age of the mosquito population and the rate of anthrophagy. This assumption will cause each individual to experience the same relative reduction in molecular force of infection, i.e. the number of new *P. falciparum* clones acquired over time. Consequently, model predictions are likely to underestimate the variance in the reduction of within-host parasite genetic diversity resulting from vector based interventions. This effect would lead to a decrease in the statistical power of the genetic metrics considered and subsequently the sample sizes presented within the power analysis are likely on the lower end of the sample sizes required for a given predictive power.

Thirdly, while the developed statistical model provided accurate estimates of malaria prevalence overall for the five regions, the prediction for Jinja was noticeably worse, which reflects the high COI observed in that region given its comparatively low prevalence. While we were able to replicate the COI age relationship for this region during model parameterisation, this was largely due to the fact that the historic prevalence for the region was much higher. For this reason, the model predicts that individuals in the region will have higher acquired immunity and will subsequently be able to harbour more infections before developing a fever and potentially being treated and thus clearing infections. The developed statistical model, however, did not include any covariates for historic prevalence or genetic diversity. Subsequently, predictions made by this model largely reflect the mean diversity expected for a given prevalence and will suffer when making predictions for regions that have experienced a recent and large decline in prevalence. Recent declines in prevalence will cause individuals in the region to possess higher immunity than predicted based solely on the region’s current prevalence, which has been shown to manifest in clear patterns in the size of the submicroscopic reservoir^34^. From a genetic perspective, increased immunity may either lead to a reduction in within-host genetic diversity due to more infections being suppressed. Alternatively, increased immunity may increase within-host genetic diversity if the higher immunity decreases the frequency with which people develop clinical symptoms, which in turn reduces the likelihood that an individual has recently been treated and subsequently has cleared all parasite strains. The latter may be a possible explanation for the comparatively high COI observed in the Walukuba study site in Uganda compared to its malaria prevalence. Consequently, as more genetic data is collected over time we will be able to extend the methods presented here to better handle recent changes in prevalence and incorporate historic measures of genetic diversity for more accurate predictions of malaria prevalence. Alternatively, the modelling framework presented here could be extended to incorporate alternative data sources, such as longitudinal measures of clinical incidence from passive surveillance.

In our model we have only considered neutral genetic markers that are unlinked. While these loci are informative for capturing standing genetic diversity, we have not considered how selective events may shape the genetic diversity. For example, if drug resistance were to spread quickly through an area it is likely that this would cause a decrease in genetic diversity in neighbouring regions^35^. However, the precise impact that this will have on the metrics explored in this study will depend on both how quickly recombination will result in linkage disequilibrium decay and the strength of the selective sweep. Although these were not assessed in this paper, it would be possible to adapt our model to consider loci under selection and simulate how known factors that affect the speed of selection, such as transmission intensity, importation of resistance, treatment rates and the metabolic costs associated with resistance, impact genetic metrics. Lastly, the model could also be extended to better capture importation and spatial dynamics. The current model employs a continent-island assumption, where the genotypes of imported parasites are drawn from a population with a fixed population-level allele frequency. This could be extended to consider populations within a metapopulation, where importations are sampled from connected populations. This would have the benefit of better capturing dynamics between different populations and could incorporate different data sources such as mobile phone records and travel surveys, which have been used to give a greater resolution to the spatial dynamics of malaria transmission^36,37^.

The 2018 world malaria report shows that the reductions in the global burden of malaria made since 2000 may be stalling, with 2 million more cases of malaria estimated in 2017 compared to 2016^38^. These declines have necessitated the development of new tools to enhance current surveillance efforts. In this study, we have shown that that malaria genetic metrics could provide an additional toolkit for operational surveillance. In particular, a combination of metrics focussed on the complexity of infections, the frequency and uniqueness of genotyped barcodes and measures of identity-by-descent could be used for inferring the prevalence of malaria across the current range of malaria prevalence. It is important to highlight that there is still a need to understand the cost-effectiveness of these tools compared to current surveillance methods. In many endemic areas, clinical incidence data provides a temporally and spatially rich measure of malaria transmission. However, it is reliant on the accuracy of estimates of the population size. In situations where this is not possible, such as migratory populations and clinics with unknown health facility catchment areas. Consequently, there may be a niche for parasite genetics to complement measures of malaria incidence in as well as in areas in which the spatial coverage of surveillance data is poor. It is hoped that these findings, in particular the importance of sample metadata and quantifying the contribution of cotransmission and superinfection events have in shaping genetic diversity, can guide future efforts by the wider community for utilising malaria genotyping for epidemiological surveillance.

## Methods

### *P. falciparum* transmission model

An individual-level stochastic model was developed to simulate the transmission dynamics of *Plasmodium falciparum*. The model is based upon previous modelling efforts^25,39–41^, however with extensions to now include individual mosquitoes as well as humans, and with parasites now modelled as discrete populations associated with individual infection events. Each parasite population is identified by a 24-SNP barcode, with sexual stages represented by two barcodes to characterise the female and male gametes within the vector and allow recombination to be explicitly modelled. An overview of the original model is given here before describing the changes made to the model, with the full methods detailed in the Supplementary Methods.

People exist in one of six infection states, with individuals beginning life susceptible to infection. At birth, individuals possess a level of maternal immunity that decays exponentially over the first 6 months. Each day individuals experience a force of infection that depends on their level of immunity, biting rate and the abundance of infectious mosquitoes. Infected individuals, after a 12-day latent period, develop either clinical disease or asymptomatic infection dependent on their level of acquired immunity from previous infections. Individuals that develop disease have a fixed probability of being effectively treated. Treated individuals enter a protective state of prophylaxis, before returning to susceptible. Individuals that did not receive treatment recover to a state of asymptomatic infection. Asymptomatic individuals progress to a subpatent infection, before clearing infection and returning to susceptible. All infected individuals that are not in the prophylactic state are also susceptible to superinfection.

The adult stage of mosquito development is modelled individually, with adult mosquitoes beginning life susceptible to infection. Mosquitoes seek a blood meal on the same day they are born and every 3 days after that until they die. Infected mosquitoes pass through a latent infection stage that lasts 10 days before becoming onwardly infectious to humans. The introduction of vector based interventions leads to a decrease in the average age of the mosquito population throughout the duration of the intervention due to the increased mortality rate. A decrease in anthrophagy is also observed reflecting mosquitoes that are repelled as a result of interventions but do not die. The daily rate of change to these parameters in response to insecticide treated nets (ITN) and indoor residual spraying (IRS) is calculated using an equivalent deterministic version of the earlier model that included interventions 25, before being introduced as a time-dependent variable within the stochastic model.

### Parasite genetics

Parasites are modelled as discrete populations that result from an infection event associated with a mosquito or a human (Supplementary Methods for full description). Each asexual parasite is characterised by one genetic barcode, which contains information relating to 24-SNPs distributed across the parasite genome. In simulations modelling identity-by-descent (IBD), the barcode is modified to contain 24 integer values that uniquely index an individual in the starting population, enabling ancestry to be tracked over time and hence IBD rather than identity-by-state (IBS) to be modelled. Sexual stages of the parasite lifecycle within the mosquito are represented by both a female and male barcode, thus defining the range of recombinants that could be produced. During a successful human to mosquito infection event, multiple oocysts may develop within the mosquito. The number of oocysts formed is drawn from a zero-truncated negative binomial distribution with mean equal to 2.5 and shape equal to 1 (95% quantile: 1-9)^42–44^, with required gametocytes sampled from the human according to the relative parasitemias of the gametocytogenic strains. During a successful mosquito to human transmission event, multiple sporozoites may be onwardly transmitted, with the genotypes the result of recombination events from ruptured oocysts. Recombination is simulated at this stage, and generated recombinants stored within the mosquito and associated with the oocyst from which it originated. The number of sporozoites passed on is drawn from a zero truncated geometric distribution with a mean of 10 (95% quantile: 1-29)^31,45^, with the percentage of sporozoites that survive estimated within model fitting.

### Model Fitting

Our extensions to the transmission model introduced a new parameter, *ζ*, which determines the percentage of the total sporozoites passed on within a feeding event that survive to yield a blood-stage infection and subsequently produce gametocytes. To fit this parameter we compared the model-predicted relationship between the complexity of infection (COI) and age utilising previously SNP genotyped samples from five sites across Kenya^46^ and Uganda^26^, collected between 2008-2010 and 2012-2013 respectively. In brief, dried blood spots were collected, and samples taken from individuals with evidence of asexual parasitemia by microscopy were selected for Sequenom SNP genotyping. Genotyping was conducted using the Sequenom MassARRAY iPLEX platform, yielding minor and major allele frequencies.

We applied *THE REAL McCOIL* proportional method to the SNP genotyped samples to estimate each individual’s COI^26^. Samples were filtered first by excluding loci with more than 20% missing samples, followed by samples with more than 25% missing loci. We performed thirty MCMC repetitions for each sample, with a burn-in period of 10^4^ iterations followed by 10^6^ sampling iterations, with genotyping measurement error estimated along with COI and allele frequencies, and a maximum observable COI equal to 25. Default priors were assigned for each parameter, and we used standard methodology to confirm convergence between chains^47^.

The observed relationship between COI and age was compared to the model-predicted relationship for each administrative region studied. The model-predicted relationship was generated by conducting simulations calibrated to estimates of the administrative malaria prevalence from 2000 to 2015^48^, exploring 50 values of *ζ* between 0.5% - 50%. For each region, 10 stochastic realisations of 100,000 individuals were simulated with a burn-in period of 50 years to ensure both an epidemiological and genetic equilibrium was reached by year 2000. For each of the five administrative regions of interest, we incorporate the historical scale up of insecticide treated nets and indoor residual spraying between 2000 and 2015, using data previously collated for the World Malaria Report 49, and estimates for the coverage of treatment modelled using DHS and MICS survey data^50^. Seasonality for each region was included by altering the total number of mosquitoes using annually fluctuating seasonal curves fitted to daily rainfall data from 2002 to 2009^51^. Lastly, we introduced rates of importation of infections that were calculated for each year between 2000 and 2015 using a fitted gravity model of human mobility^52^. These sources represent infections acquired from individuals travelling out of the region and returning with an infection, and also mosquitoes being infected by individuals travelling from outside into to the region of interest.

We calculated the “distance” between our model predictions and the observed data using the Kullback-Leibler (KL) divergence^53^. Using an individual’s age and estimated COI, the distance between the observed and predicted distributions of COI for each age is given by:

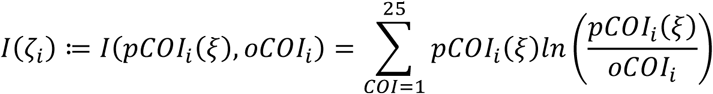

where *oCOi*_*i*_ is the observed distribution of COI at age *i* and *pCOi*_*i*_ (*ζ*) is one realisation of the model-predicted distribution of COI at age *i* for a given frequency of successful sporozoites *ζ* (with only parasites that would have been detected by PCR being assumed to be detected by SNP genotyping). The total distance for a given value of *ζ* is subsequently given by:

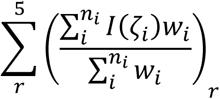

where *w*_*i*_ is the weight for age *i*, and *n*_*i*_ is the total number of unique sampled ages in administrative region *r*. This can be interpreted as the sum of the weighted KL divergence means within a region, with weights equal to the number of observations at each age. Each region thus contributes equally to the total distance, despite the difference in the number of individuals in each region.

Further model fit validation was conducted by incorporating a comparatively larger collection of estimates of the COI estimated using *msp2* genotyping, which is more commonly referred to as multiplicity of infection (MOI). *msp2* genotyping is known to known to underestimate COI in individuals with very high COIs, with COIs > 7 difficult to observe. Consequently, to distinguish these estimates we refer to these as *msp2* COI. We compiled *P. falciparum* malaria MOI data where there were estimates of both the malaria prevalence and the MOI of study participants. This was conducted by updating a previous review^16^, using the same search terms of “falciparum multiplicity infection prevalence msp2”. Analogous relationships were predicted using the fitted model, with the model predicted *msp2* COI estimated by assuming that any individual with a model predicted COI greater than 7 results in an *msp2* COI of 7, which reflects the limits of resolution when using *msp2* genotyping^54^.

### Contribution of superinfection and cotransmission events towards within-host genetic diversity

The parameterised model was used to characterise the relative contribution of cotransmission events and superinfection events towards within-host parasite genetic diversity. Ten stochastic realisations of 100,000 individuals were simulated for 50 years at 15 different transmission intensities. The proportion of highly identical parasite strains (>50% of loci are IBD in pairwise comparison) within simulations was recorded and used to estimate the proportion of within-host genetic diversity that is due to cotransmission events rather than superinfection.

### Impact of changes in transmission intensity upon measures of parasite genetic diversity

The effect of declines in transmission intensity on four measures of within-host genetic diversity was explored. The four measures considered were: 1) the mean COI, 2) the percentage of polygenomic infections (% Polygenomic), 3) the percentage of unique barcode genotypes (% Unique), and 4) a newly defined metric, the coefficient of uniqueness (COU), which is given by:

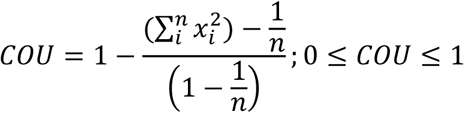

where *x*_*i*_ is the frequency at which barcode *i* occurs within a sample of size*n. COU* = 0 when all barcodes within a sample are identical, and *COU* = 1 when all barcodes within a sample are unique.

Ten stochastic realisations of 100,000 individuals were simulated for 50 years with an initial parasite prevalence measured by PCR equal to ∼70% and a fixed importation rate to ensure both a genetic and epidemiological equilibrium. Once at equilibrium, three differing levels of intervention scale-up (low, medium, high) were introduced that lead to an absolute reduction in parasite prevalence from 70% to 45%, 20% and 5% after 10 years. The scale-up of interventions resulted in an increase in the coverage of ITNs (maximum after 10 years: 30%, 60%, and 90%), IRS (maximum after 10 years: 20%, 40% and 60%) and treatment (maximum after 10 years: 15%, 30%, 45%). For all simulations, the monthly mean for each genetic marker was recorded for the whole population as well as within three age ranges (0-5 years old, 5-15 years old and over 15 years old), and within individuals who were asymptomatic or symptomatic at the time of sample collection.

An identical analysis was conducted at a lower starting prevalence, with maximum reductions in parasite prevalence by PCR from 35% to 20%, 2% and ∼0% after 10 years, in order to assess the change in two measures of identity-by-descent (IBD), pIBD and iIBD. The population mean IBD (pIBD) we define as the mean number of loci in pairwise comparisons between samples that are identical across all loci in terms of their 24-locus identity barcode (focusing on genotypes that could be detected by microscopy only), i.e. it is the mean proportion of shared ancestry between samples. The individual mean IBD (iIBD) is the mean number of identical loci of the 24-locus identity barcode within individuals who are polygenomically infected. If all sampled individuals are monogenomic, then iIBD is set equal to 1.

### Statistical power analysis of parasite genetic measures

To evaluate the utility of the considered measures of parasite genetic diversity, we conducted an analysis to characterise the predictive power of each metric for detecting changes in transmission intensity, and their sensitivities to the sample size chosen. In an analogous design to earlier simulations, we measured sample mean measures of the COI, % Polygenomic, % Unique, COU, iIBD and pIBD at yearly intervals for the first five years after the initiation of the ten-year scale-up of interventions.

Sensitivity to the sample size of each metric was assessed by sequentially sampling subsets of the data and comparing the mean difference in metrics. Sample sizes between 10 and 600 individuals were explored, with 100 samples drawn from a stochastic realisation at years 0, 1, 2, 3, 4 and 5, and comparisons made between years 1-5 and year 0, i.e. 0-1, 0-2, … 0-5. All samples were collected from individuals aged between 5-15 years old. One-tailed Monte Carlo p-values were generated for each subsample by 100 permutations of the years that samples were collected from. The power of each metric was defined as the proportion of subsamples for which 95% of the permuted mean differences were greater or less than the observed mean difference, with the direction of the tail dependent on whether the metric is expected to decrease or increase respectively in response to a decrease in transmission intensity. The overall power for each metric was calculated as the mean power of ten stochastic realisations, and repeated at two different starting parasite prevalence by PCR (∼60% and ∼30%). Metrics based on comparisons of IBD were only assessed for the lowest starting parasite prevalence. The performance of each metric was also explored under the assumption that it was not possible to phase all genotypes within the samples collected, and that only the dominant genotype was able to be called.

### Statistical modelling of the predictive performance of malaria genomics for surveillance

A statistical model was constructed to predict malaria prevalence using the genomic metrics explored thus far, with three different assumptions about the availability of patient metadata (no metadata, patient age only, and both patient age and symptomatic status of infection). To assess the utility of such a model for surveillance, samples of 200 individuals were taken from a range of simulations that span the transmission, seasonality and intervention coverage range seen in sub-Saharan Africa. We used the sampled mean measures of the genomic metrics discussed, and where available summaries of the age and clinical status of samples to create our model simulated datasets. 25% of simulated datasets were held back for out-of-sample testing. Three different statistical models (gradient boosted trees, elastic net regression model and random forests) were fit to the model simulated data. The predictions of these level 1 models were subsequently used to train an ensemble model using a linear optimisation based on the root mean squared error (RMSE) of the level 1 models. When training both the level 1 models and the ensemble, K-fold cross validation sets were performed 25 times and subsequently averaged to reduce any bias from the cross validation set chosen. The averaged cross validation results were used to assess the performance of the ensemble model on the testing dataset by comparing the RMSE, mean absolute error (MAE) and the correlation under the different assumptions about the availability of patient metadata. The predictors of the ensemble model were assessed for their contribution to the overall model performance. Variable importance was calculated for each level 1 model, before reporting their overall importance as the weighted mean importance, with the weight equal to the level 1 model weights in the ensemble model. Lastly, the trained ensemble model was used to predict the prevalence of malaria for the study sites considered within Uganda and Kenya.

## Acknowledgements

OJW and JH acknowledge funding from Wellcome Trust PhD Studentships (109312/Z/15/Z and 105272/Z/14/Z). HJTU, LCO and ACG acknowledge grant support from the Bill and Melinda Gates Foundation. LCO also acknowledges funding from a UK Royal Society Dorothy Hodgkin fellowship. LCO and ACG acknowledge Centre support from the Medical Research Council and Department for International Development. Kenyan school surveys and sample collections were funded by the Division of Malaria Control, Ministry of Public Health and Sanitation through a grant from DFID through the WHO Kenya Country Office. RWS acknowledges funded as a Principal Wellcome Fellow (103602 & 212176). H-HC was funded by the National Institute of General Medical Sciences (U54GM088558). RV is funded by a Skills Development Fellowship: this award is jointly funded by the UK Medical Research Council (MRC) and the UK Department for International Development (DFID) under the MRC/DFID Concordat agreement and is also part of the EDCTP2 programme supported by the European Union.

## Author Contributions

OJW drafted the paper. OJW, LO, ACG and RV conceptualized the study. OJW developed software with additional input from JH, HCS, HJTU and RV. OJW and RV conducted data analysis with additional input from H-HC, LCO and ACG. IO, PB, RWS, AMN, KR, CH, JIN, BG were involved in data collection. All authors contributed to interpretation of the analyses and revised the draft paper.

## Competing interests

The authors declare no competing interests.

## Supplementary Material

**Supplementary Figure 1:**
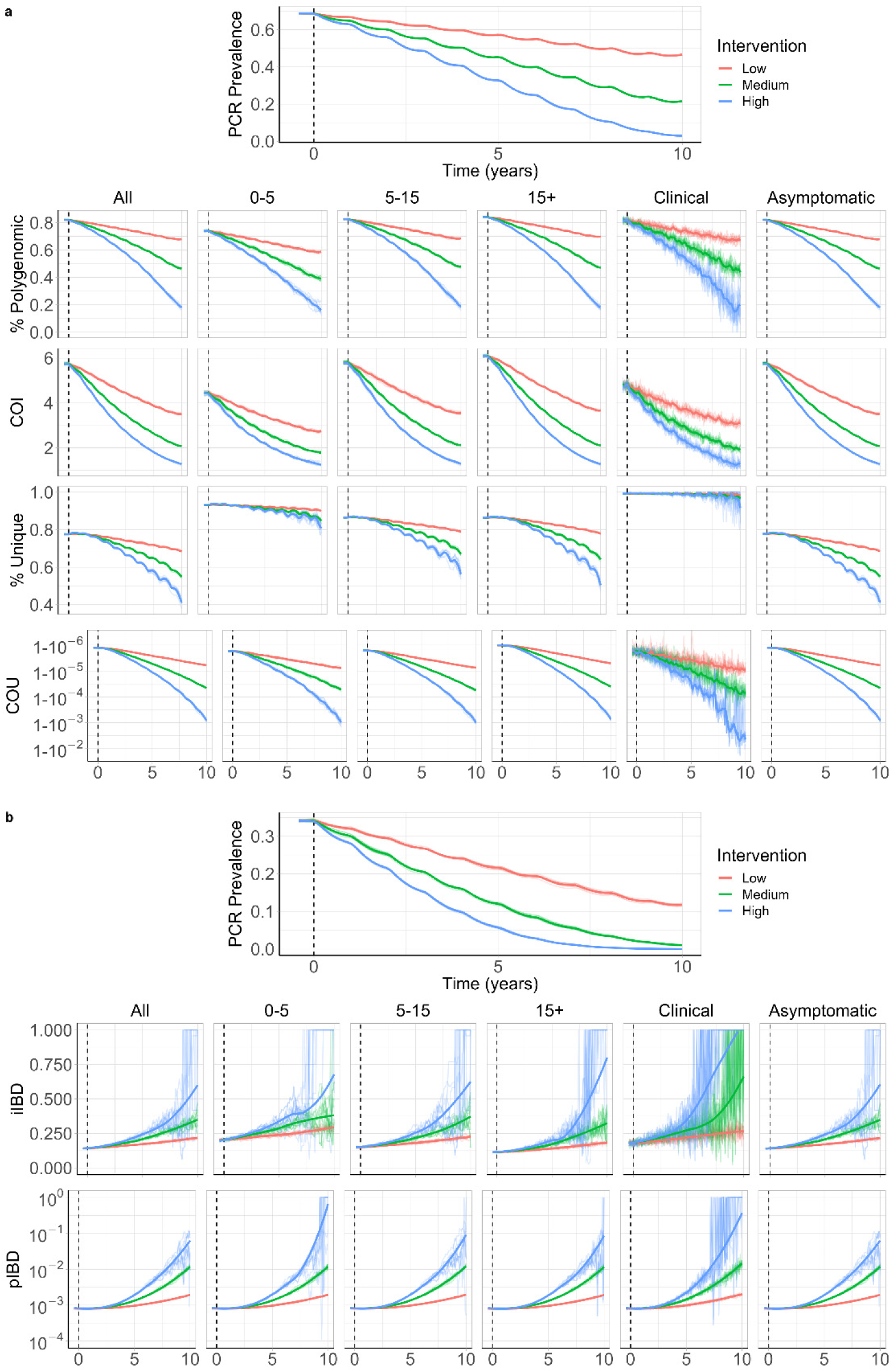
Age and sampling dependent impact of changes in transmission intensity upon genetic metrics of transmission intensity. In **a)** the top plot shows the change in PCR prevalence after the introduction of 3 different levels of intervention scale up, with both the 10 individual stochastic realisations and the mean local regression smoothed relationship shown. The following four rows show the population mean percentage of the population that are polygenomically infected, the complexity of infection (COI), the percentage of samples that are genotypically unique (% Unique) and the coefficient of uniqueness (COU) for the prevalence declines seen in the first row. The metrics are stratified into columns by the sampling scheme chosen. In **b)** the top plot shows the change in PCR prevalence, which reaches <1% in the highest intervention arm. The following rows show the within host identity-by-descent (iIBD) mean across the 24 identity loci considered, and the population mean pairwise measure of IBD (pIBD). In both the same sampling stratification is used as in **a)**. In all plots the vertical dashed black line shows the time from which the scale up of interventions starts (Time = 0 years).

**Supplementary Figure 2:**
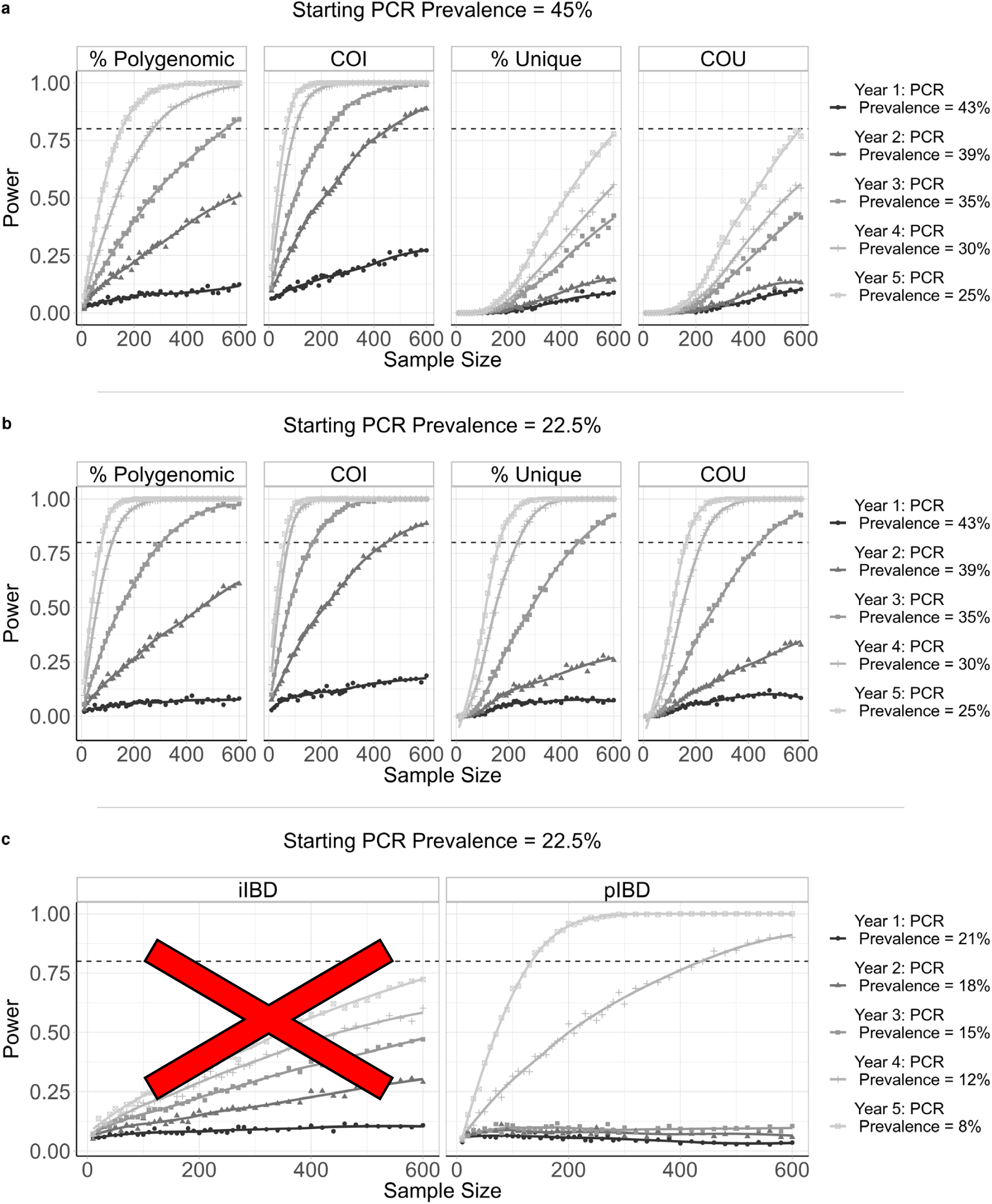
Predictive power of six metrics of parasite genetic diversity with respect to sample size under the assumptions that samples are unable to be phased. The same methods as those detailed in the main text were used, with the only difference being that samples could not be phased and only the major haplotype could be called for an individual. iIBD is unable to be measured if samples cannot be phased and is subsequently crossed out. For pIBD, % Unique and COU it was assumed that the highest parasitaemia barcode was detected from each polygenomically infected individual. Lastly, there was no assumed difference in the ability to detect polygenomic samples or estimate the COI with unphased samples.

**Supplementary Figure 3:**
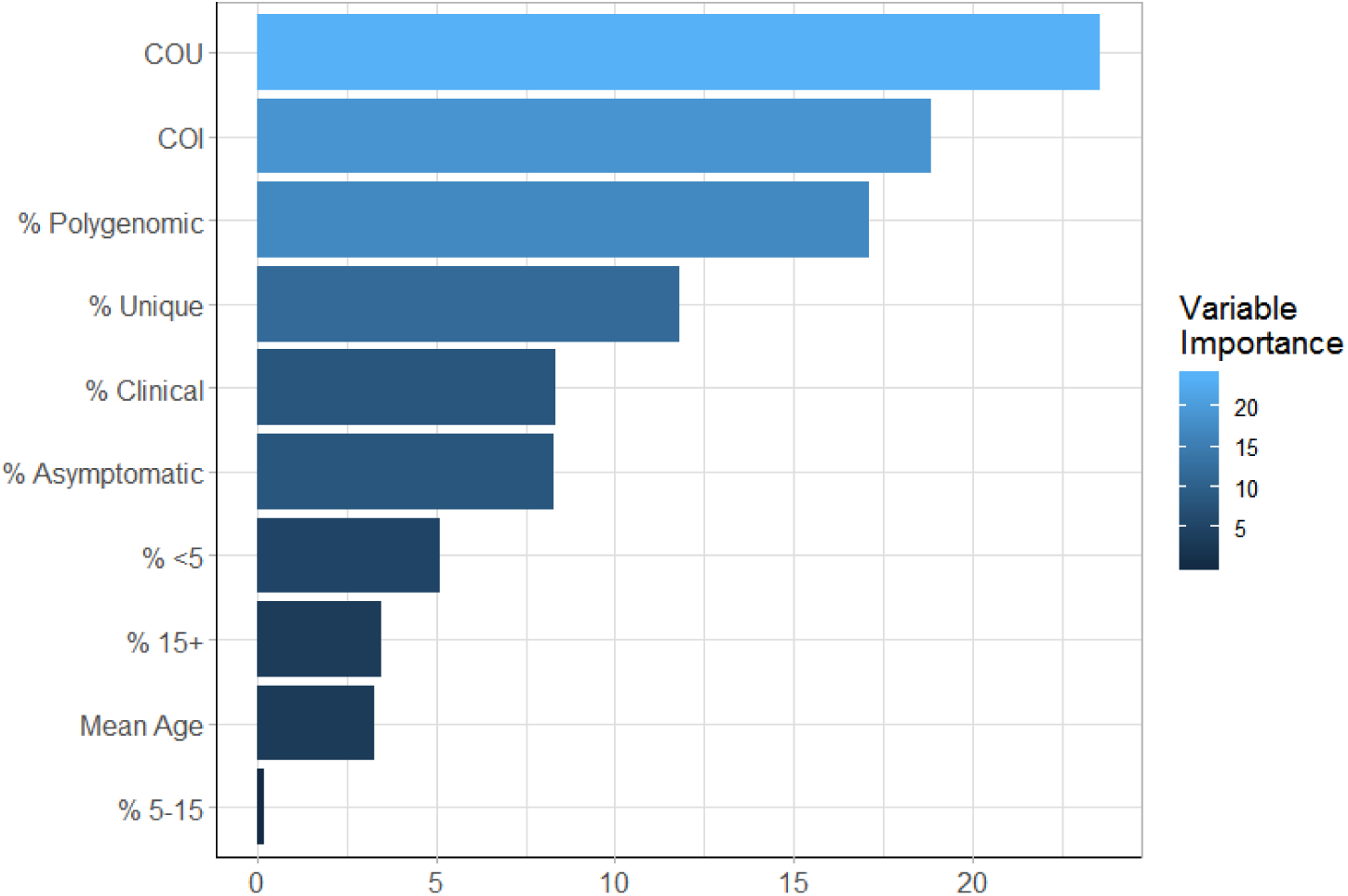
Mean Importance of each predictor variable within the trained ensemble model for predicting malarial prevalence. The newly defined measure, the coefficient of uniqueness (COU), was observed to be the most important metric, with the six metadata variables (age and clinical status) being the least important. They do, however, contribute 28% of the total model importance, which highlights why the inclusion of this metadata resulted in better model predictions.

**Supplementary Figure 4:**
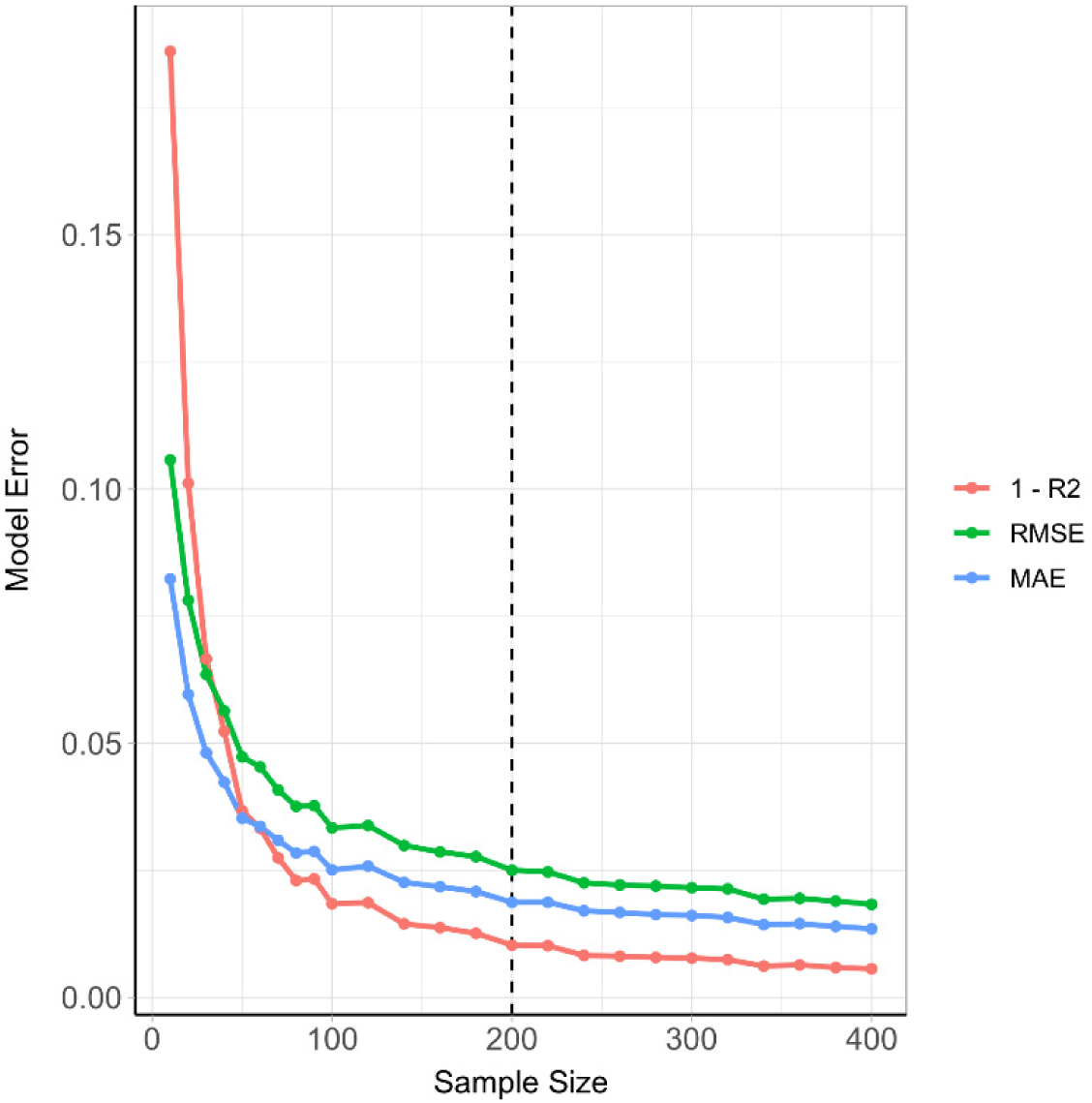
The predictive performance of the ensemble model under different assumed sample sizes. Measures of the model error, root mean squared error (RMSE) and root mean error (MAE) as well as *1* − *R*^*2*^ are shown for sample sizes between 10 and 400. Model performance improves quickly over sample size ranges between 10 and 100, before slowing, with only very modest increases seen in model performance for sample sizes larger than 200.

**Supplementary Figure 5:**
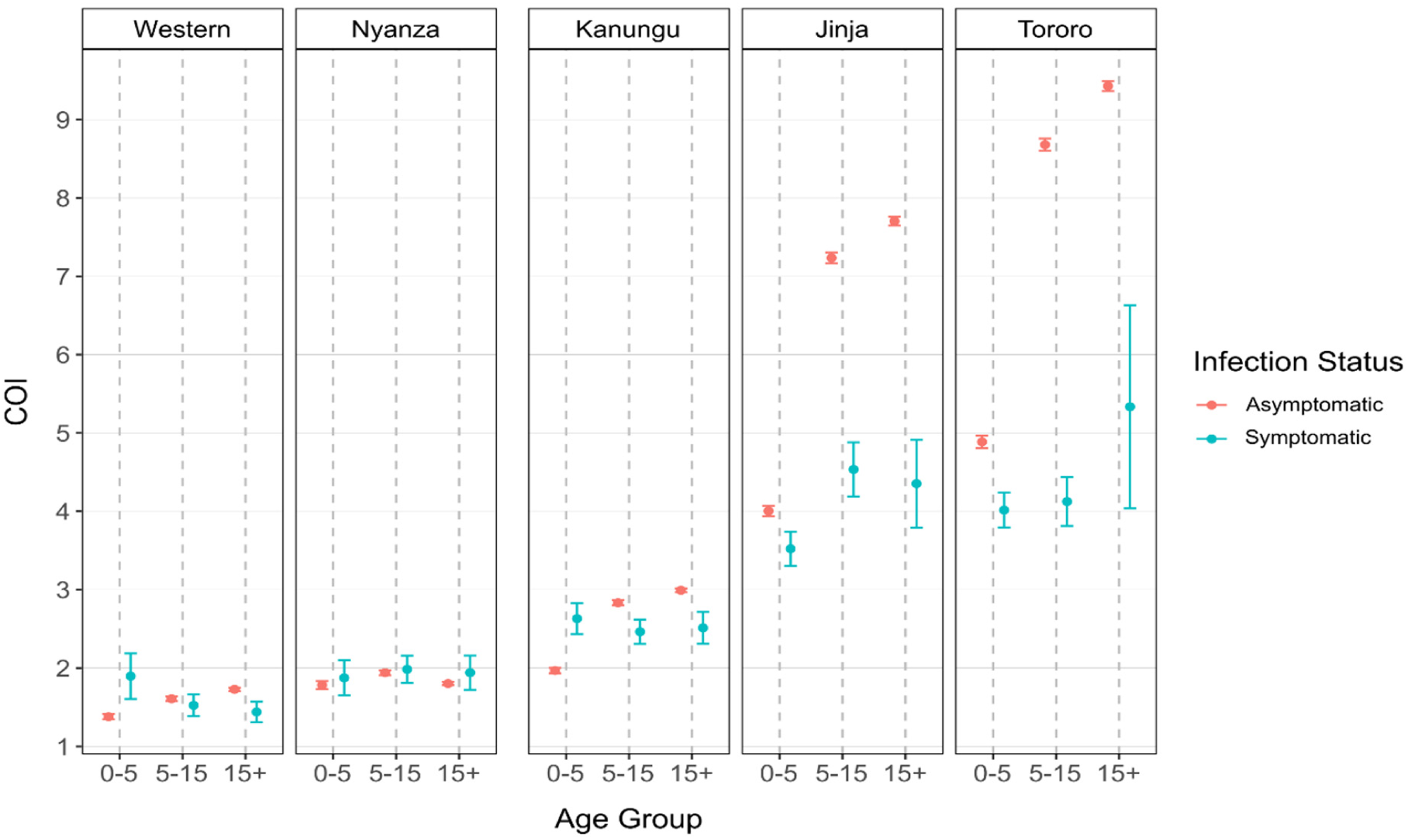
Age and symptomatic status stratified COI from model predictions during the model fitting. Each plot shows the mean COI and 95% confidence interval for the study sites used in the model fitting. COI is stratified by age group and symptomatic status, showing that on the whole COI is higher in asymptomatic individuals, however, in lower transmission areas COI is higher in symptomatic young children.

**Supplementary Figure 6:**
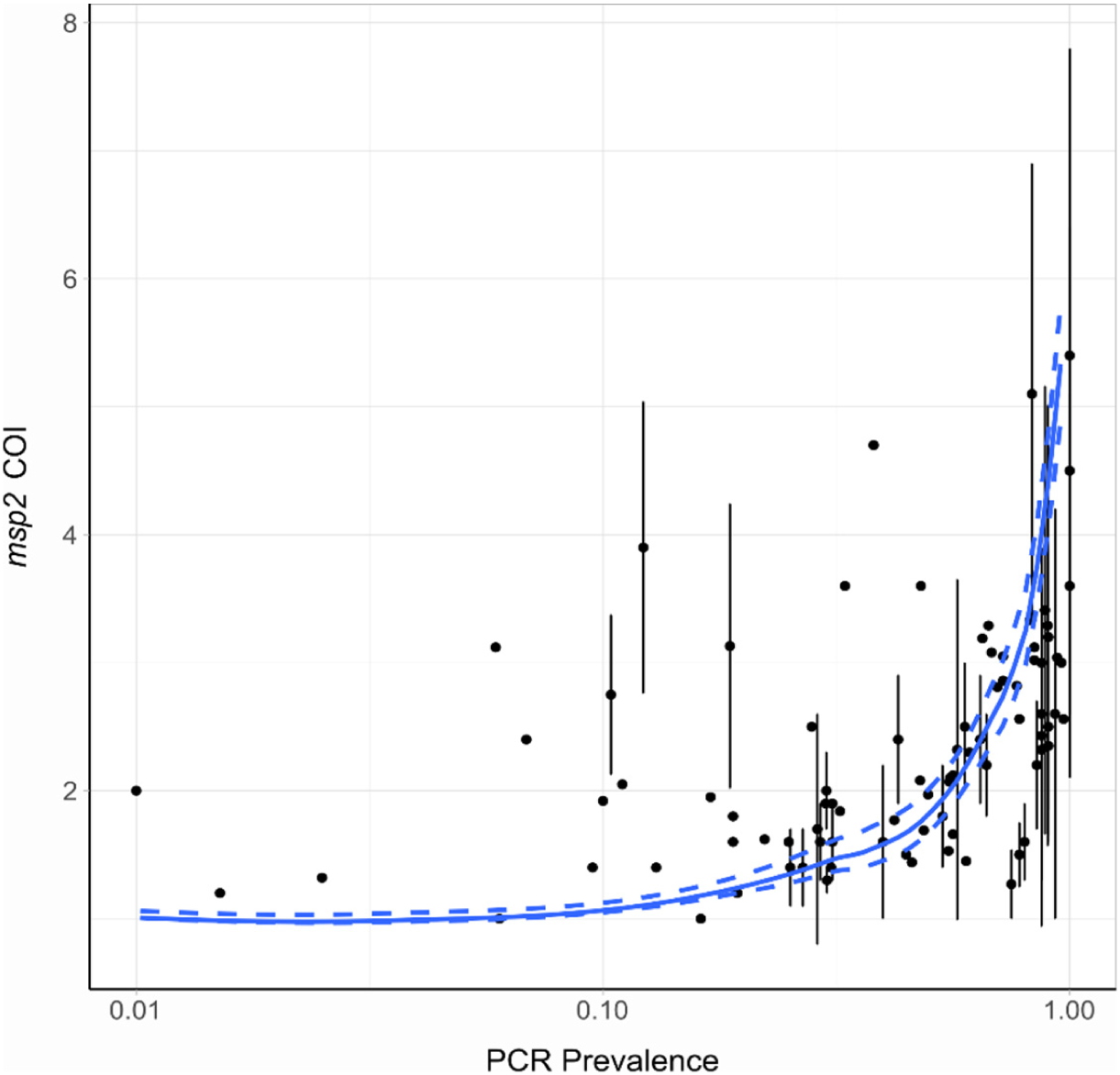
Model predicted relationship *msp2* COI and PCR prevalence. The blue solid line shows the relationship for the fitted value of ζ equal to 0.20. The dashed lines above and below this in blue show the relationship for values of ζ equal to 0.29 and 0.10 respectively. The point-ranges in black show the observed values of COI by *msp2* genotyping from the literature review.

**Supplementary Figure 7:**
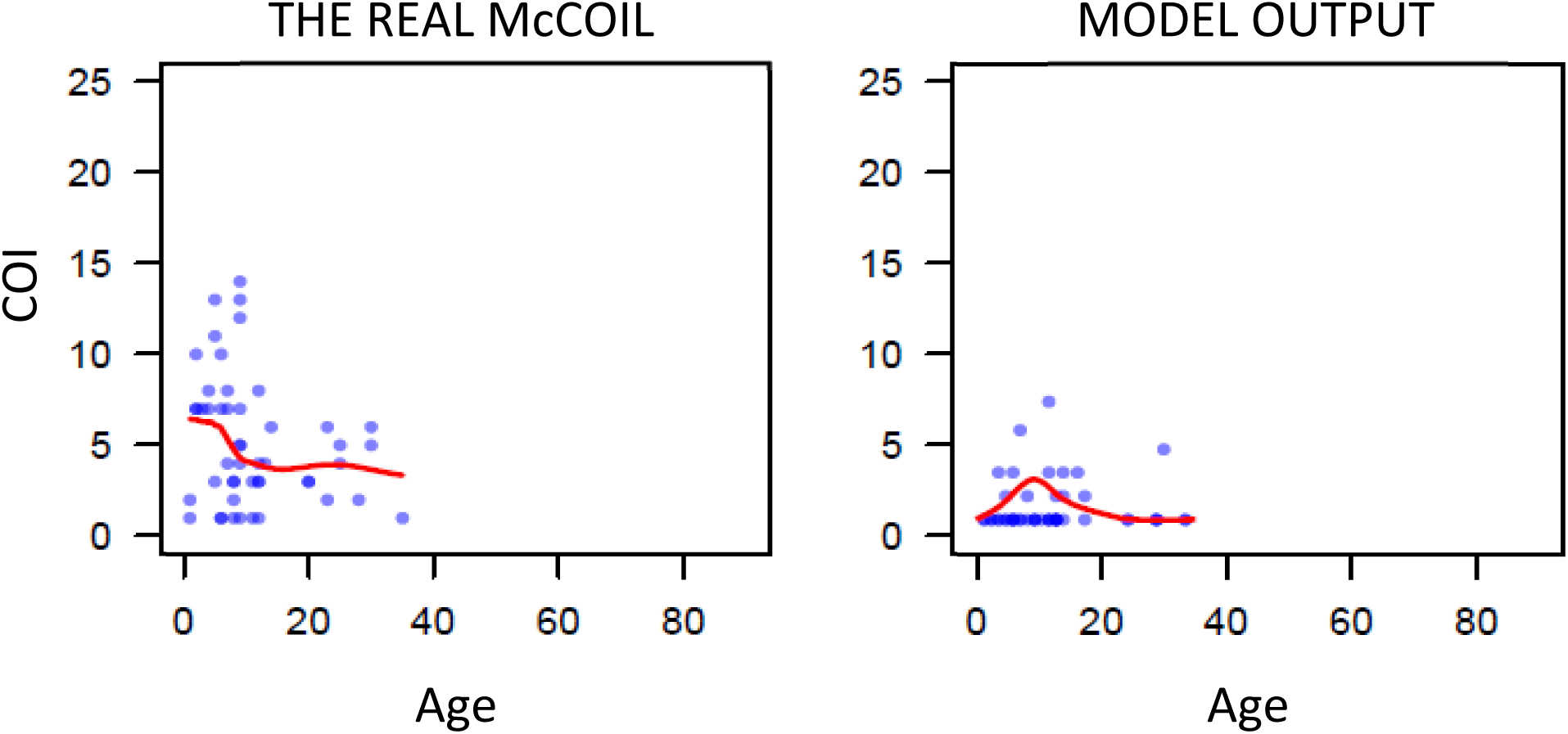
The fitted model-predicted relationship between COI and age for Walukuba, if the prevalence simulated was assumed to be equal to the prevalence within the sub-county surveyed, rather than the prevalence for the administrative region. Model fitting conducted in Figure 1 in the main text used the administrative region prevalence as estimated by the Malaria Atlas Project, which resulted in good agreement between COI and prevalence.

**Supplementary Table 1:** Statistical Model Performance.

| Meta Data | Model | RMSE* | MAE* | R<sup>2</sup>* |
| --- | --- | --- | --- | --- |
| None | Elastic Net | 0.0276 (157.71%) | 0.0225 (173.08%) | 0.9935 (99.61%) |
| None | Gradient Boosted Trees | 0.0214 (122.29%) | 0.0159 (122.31%) | 0.9961 (99.87%) |
| None | Random Forest | 0.0211 (120.57%) | 0.0151 (116.15%) | 0.9962 (99.88%) |
| None | Weighted Mean Ensemble | 0.0204 (116.57%) | 0.0151 (116.15%) | 0.9965 (99.91%) |
| Age | Elastic Net | 0.0311 (177.71%) | 0.0245 (188.46%) | 0.9921 (99.47%) |
| Age | Gradient Boosted Trees | 0.02 (114.29%) | 0.0152 (116.92%) | 0.9967 (99.93%) |
| Age | Random Forest | 0.0197 (112.57%) | 0.0143 (110%) | 0.9968 (99.94%) |
| Age | Weighted Mean Ensemble | 0.0195 (111.43%) | 0.0144 (110.77%) | 0.9969 (99.95%) |
| Age and Clinical Status | Elastic Net | 0.0278 (158.86%) | 0.0219 (168.46%) | 0.9934 (99.6%) |
| Age and Clinical Status | Gradient Boosted Trees | 0.0178 (101.71%) | 0.0141 (108.46%) | 0.9974 (100%) |
| Age and Clinical Status | Random Forest | 0.0178 (101.71%) | 0.013 (100%) | 0.9973 (99.99%) |
| Age and Clinical Status | Weighted Mean Ensemble | 0.0175 (100%) | 0.013 (100%) | 0.9974 (100%) |

\* Absolute value (% relative to best performing model)

## Supplementary Methods

### *P. falciparum* Transmission Model

An individual-based stochastic model with a fixed daily time step was developed to simulate the transmission dynamics of *Plasmodium falciparum*. Both the human and adult mosquito stages are modelled at an individual level, whereas parasites are modelled as discrete populations with each population relating to an infection event. The human transmission model is based upon previous modelling efforts^1–4^, which is described in its deterministic framework first, before detailing the human acquisition of immunity and the full set of equations detailing its stochastic implementation. The deterministic model described within the methods has been included as its equilibrium solution is used for model initialisation. Additionally, we developed a deterministic version of the earlier 2016 Griffin et al. model^1^ that incorporates interventions, which is used to indirectly incorporate the effects of intervention strategies as these are not modelled explicitly within the individual model. (The deterministic implementation of interventions has not been included within the deterministic model described below to ensure clarity related to our indirect handling of interventions).

We continue to describe the mosquito transmission model, which is again based on earlier modelling efforts^1–4^, before describing the stochastic equations detailing the new implementation of the adult mosquito stage at an individual-based level. Extensions detailing how the parasite populations are incorporated follow, by first describing the genetic barcode that each parasite population possesses. We continue by describing the within host parasite populations, which includes considerations surrounding the contribution of coinfection and superinfection towards the model’s dynamics of within-host multiplicities of infection, and how these relate to the probabilistic uptake of specific gametocyte strains by mosquitoes. This is followed by detailing the within-mosquito parasite populations, which explores the derivation of the distribution describing the model-predicted oocyte intensities, and describes how recombination within the sexual stage is explicitly modelled.

#### Human transmission model

Individuals begin life susceptible to infection (state S) (Diagram 1). At birth, individuals possess a level of maternal immunity that decays exponentially over the first 6 months. Each day individual *i* is probabilistically exposed to infectious bites governed by their individual force of infection (*Λ*_*i*_). *Λ*_*i*_ is dependent on their pre-erythrocytic immunity, exposure to bites (dependent on both their age and their individual relative biting rate due to heterogeneous biting patterns by mosquitoes) and the size of the infectious mosquito population. Infected individuals, after a latent period of 12 days (*d*_*E*_), develop either clinical disease (state D) or asymptomatic infection (state A). This outcome is determined by their probability of acquiring clinical disease (*ϕ*_*i*_), which is dependent on their clinical immunity. Individuals that develop disease have a fixed probability (*f*_*T*_) of seeking treatment (state T). Treated individuals are assumed to always recover, i.e. fully-curative treatment, and then enter a protective state of prophylaxis (state P) at rate *r*_*T*,_ before returning to susceptible at rate *r*_*s*_. Individuals that did not receive treatment recover to a state of asymptomatic infection at rate *r*_*D*_. Asymptomatic individuals progress to a subpatent infection (stage U) at rate *r*_*A*_, before clearing infection and returning to susceptible at rate *r*_*U*_. Additionally, superinfection is possible for all individuals in states D, A and U. Superinfected individuals who receive treatment will move to state T. Individuals who are superinfected but do not receive treatment in response to the superinfection will either develop clinical disease, thus moving to state D, or develop an asymptomatic infection and move to state A (except for individuals who were previously in state D, who will remain in state D).

**Diagram 1:**
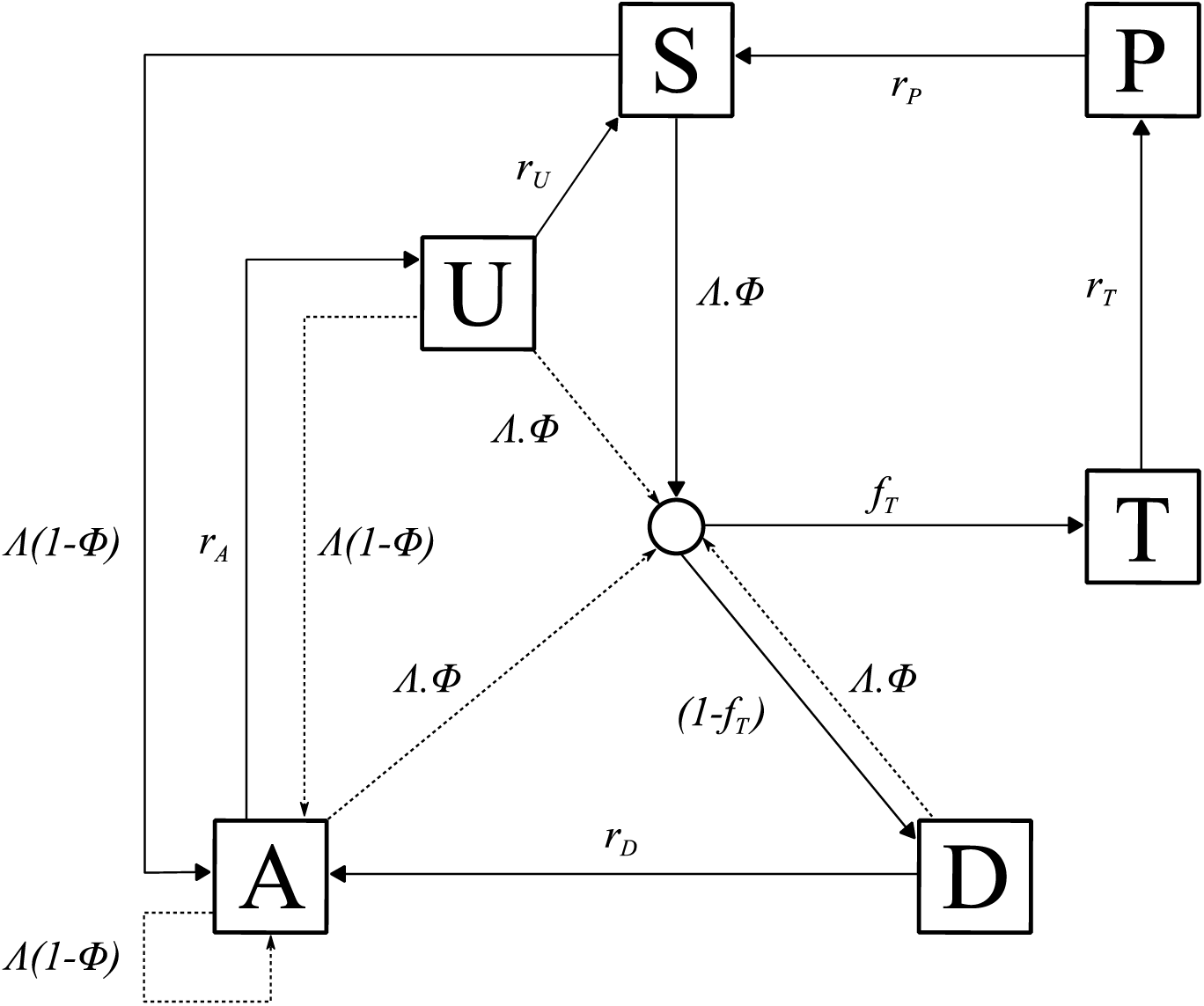
Transmission Model. Flow diagram for the human component of the transmission model, with dashed arrows indicating superinfection. S, susceptible; T, treated clinical disease; D, untreated clinical disease; P, prophylaxis; A, asymptomatic patent infection; U, asymptomatic sub-patent infection. All parameters are described and referenced within Table 1.

The movement between the human components of the transmission model is summarised with the following partial differential equations describing each compartment (*t* represents time and *a* represents age):

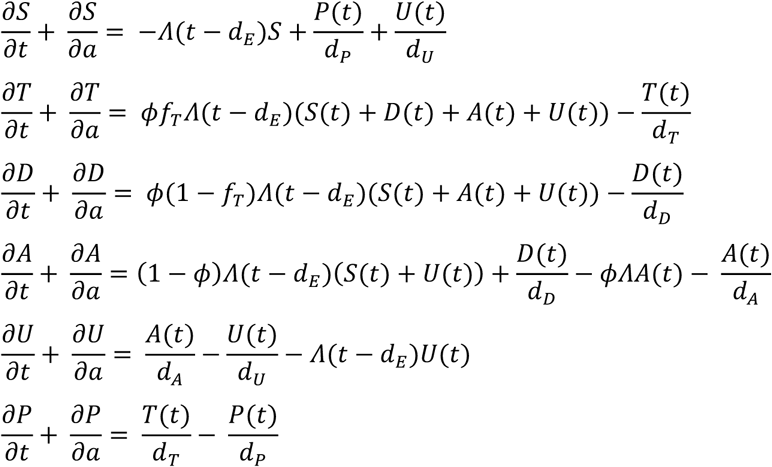

When an individual enters a new infection state a waiting time is sampled from an exponential distribution for when the individual will move out of that infection state (except when individuals move into S). With the introduction of a fixed daily time-step, the day on which an individual transitions from state X to Y occurs is given by:

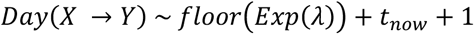

where *t*_*now*_ is the current day, i.e. the day that the individual moved into state A, and *λ* is the transition rate. The set of state transitions for individuals and their associated transition rates are given below.

| Process | Transition | Transition Rate |
| --- | --- | --- |
| Progression of untreated disease to asymptomatic infection | $D \rightarrow A$ | $r_D = \frac{1}{d_D}$ |
| Progression of asymptomatic infection to subpatent infection | $A \rightarrow U$ | $r_A = \frac{1}{d_A}$ |
| Progression of subpatent infection to susceptible | $U \rightarrow S$ | $r_U = \frac{1}{d_U}$ |
| Progression of treated disease to uninfected prophylactic period | $T \rightarrow P$ | $r_T = \frac{1}{d_T}$ |
| Progression from uninfected prophylactic period to susceptible | $P \rightarrow S$ | $r_P = \frac{1}{d_P}$ |

We assume that each person has a unique biting rate, which is the product of their relative age dependent biting rate, *ψ*_*i*_, given by

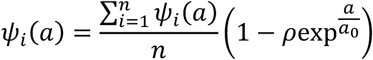

and an assumed heterogeneity in biting patterns of mosquitoes, *ζ*_*i*_, which we assume persists throughout their lifetime and is drawn from a log-normal distribution with a mean of 1,

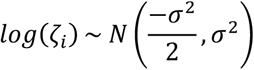

where 1 - *ρ* is the relative biting rate at birth when compared to adults and *a*_0_ represents the time-scale at which the biting rate increases with age. The product of these biting rates is subsequently used to calculate the proportion of the whole population’s bites that person *i* receives on a given day, *π*_*i*_. Their daily entomological inoculation rate (EIR), *ϵ*_*i*_, is thus calculated by multiplying by the number of infectious mosquitoes taking a blood meal from a human that day, which in turn yields their force of infection, which are given by:

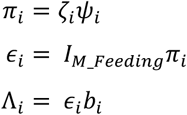

where *I*_*M_Feeding*_ is the size of the feeding infectious mosquito population, and *b*_*i*_ is the probability of infection given an infectious mosquito bite.

The inclusion of individual mosquitoes results in the following stochastic implementation of infection. On any given day the number of infectious mosquitoes taking a blood meal from a human (*I*_*M_Feeding*_) will result in the same number of infectious bites. These bites are allocated by sampling from the multinomial distribution using the conditional binomial method,^5^ where sample weights are equal to *π*_*i*_. Upon receiving an infectious bite, an individual will move to an untracked infection state, *I*, which leads to either clinical disease (*D*), treated clinical disease (*T*) or asymptomatic infection (*A*). This leads to the following transition rates related to infection below.

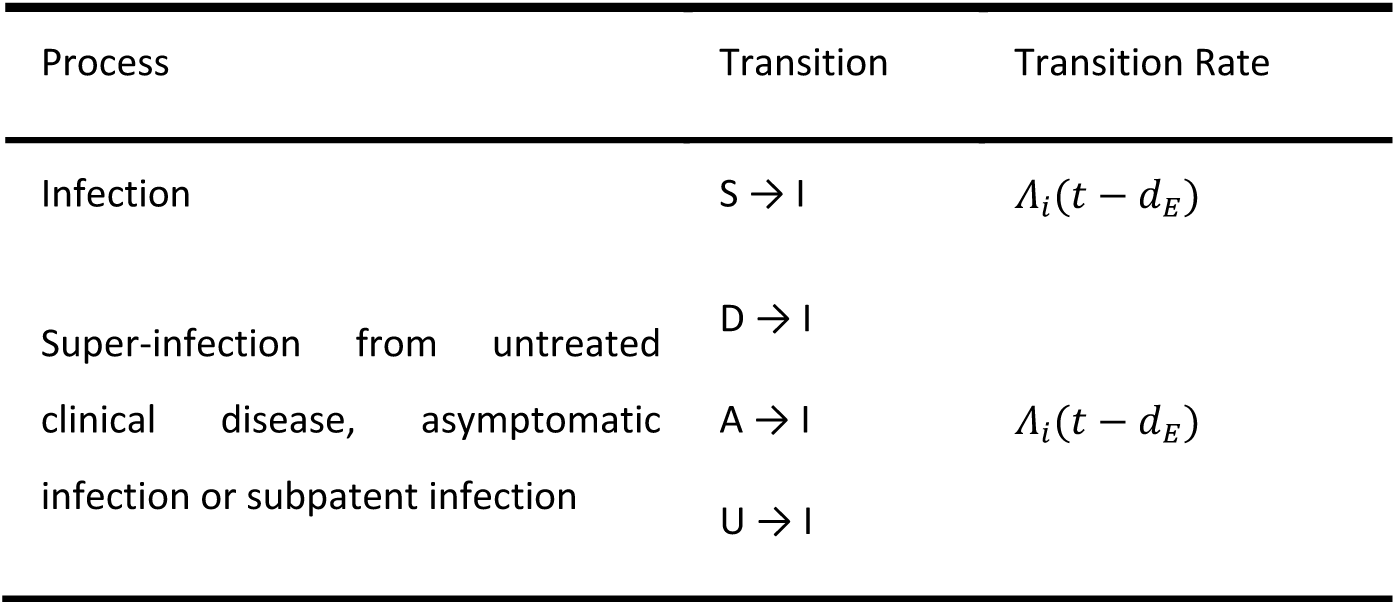

| Process | Transition | Transition Rate |
| --- | --- | --- |
| Infection | $S \rightarrow I$ | $\Lambda_i(t - d_E)$ |
| Super-infection from untreated clinical disease, asymptomatic infection or subpatent infection | $D \rightarrow I$<br>$A \rightarrow I$<br>$U \rightarrow I$ | $\Lambda_i(t - d_E)$ |

The probabilities of progressing from state *I* to *D, T* or *U* are determined an individual’s probability of clinical disease, *ϕ*_*i*_, and the treatment coverage:

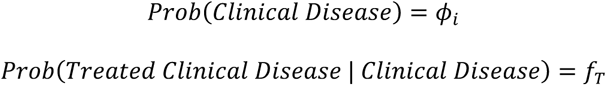

The human population was assumed to have a maximum possible age of 100 years, with an average age of 21 years within the population yielding an approximately exponential age distribution typical of sub-Saharan countries. The day on which a human dies is thus allocated at birth by sampling from an exponential distribution with a mean equal to 21 years. When an individual dies, they are replaced with a new-born individual with the same individual biting rate due to heterogeneity in biting patterns.

#### Immunity and Detection Functions

We model 3 stages at which immunity may impact transmission, as in the existing Griffin et al model:

1. Pre-erythrocytic immunity, *I*_*B*_; reduction in the probability of infection given an infectious mosquito bite.
2. Acquired and Maternal Clinical Immunity, *I*_*CA*_ and *I*_*CM*_ respectively; reduction in the probability of clinical disease given an infection due to the effects of blood stage immunity.
3. Detection immunity, *I*_*D*_; reduction in the probability of detection and a reduction in the

Maternal clinical immunity is assumed to be at birth a proportion, *P*_*M*_, of the acquired immunity of a 20 year-old and to decay at rate 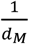. The remaining three types of immunity are described by the following partial differential equations, which describe how immunity increases due to exposure from zero at birth and decreases over time:

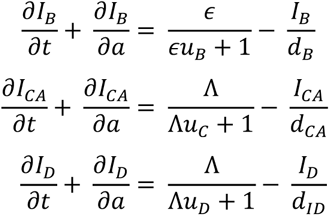

where each *u* term represents the time during which immunity cannot be boosted further after a previous boost and each *d* term represents the duration of immunity.

The probabilities of infection, detection and clinical disease are subsequently created by transforming each immunity function by Hill functions. An individual’s probability of infection, *b*_*i*_, is given by

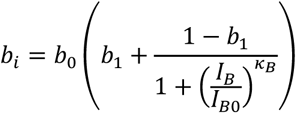

where *b*_0_ is the maximum probability due to no immunity, *b*_0_*b*_1_ is the minimum probability and *I*_*B*0_ and *κ*_*B*_ are scale and shape parameters respectively.

An individual’s probability of clinical disease, *ϕ*_*i*_, is given by

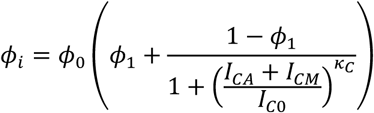

where *ϕ*_0_ is the maximum probability due to no immunity, *ϕ*_1_*ϕ*_0_ is the minimum probability and *I*_*CO*_ and *κ*_*C*_ are scale and shape parameters respectively.

An individual’s probability of being detected by microscopy when asymptomatic, *q*_*i*_, is given by

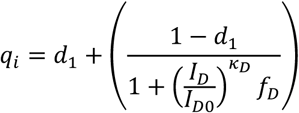

where *d*_1_ is the minimum probability due to maximum immunity, and *I*_*D*0_ and *κ*_*D*_ are scale and shape parameters respectively. *f*_*D*_ is dependent only on an individual’s age is given by

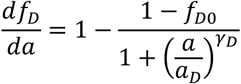

where *f*_*D*0_ represents the time-scale at which immunity changes with age, and *a*_*D*_ and *y*_*D*_ are scale and shape parameters respectively.

The probability that an infected individual infects a mosquito upon being bitten is proportional to both their infectious state and their probability of detection, with a lower probability of detection assumed to correlate with a lower parasite density. Individuals who are in state D (clinically diseased), state U (sub-patent infection) and state T (receiving treatment) contribute to an onward infection within a mosquito with probabilities *c*_*D*_, *c*_*U*_ and *c*_*T*_. In state *A*, contribution to an onward infection within a mosquito occurs with probability *c*_*A*_, and is given by *c*_*U*_ + *(c*_*D*_ − *c*_*U*_*)q*^*γI*^ where *q* is the probability of being detected by microscopy when asymptomatic, and *γ*_*I*_ is a parameter that controls how quickly infectiousness falls within the asymptomatic state.

#### Human Stochastic Model Equations

Given the definitions above, the full stochastic individual-based human component of the model can be formally described by its Kolmogorov forward equations. As before, let *i* index individuals in the population. Then the state of individual *i* at time *t* is given by *{j, k, t*_*k*_, *l, t*_*l*_, *m, t*_*m*_, *a, t}*, where *a* is age, *j* represents infection status (*S, D, A, U, T* or *P*), *k* is the level of infection-blocking immunity and *t*_*k*_ is the time at which infection blocking immunity was last boosted. Similarly, *l* and *t*_*l*_ denote the level and time of last boosting of clinical immunity, respectively, while *m* and *t*_*m*_ do likewise for parasite detection immunity. Let *δ*_*p,q*_ denote the Kronecker delta *δ*_*p,q*_ *= 1* if *p = q* and 0 otherwise) and *δ(x)* denote the Dirac delta function. Defining *P*_*i*_(*j, k, t*_*k*_, *l, t*_*l*_, *m, t*_*m*_, *a, t*) as the probability density function for individual *i* being in state *{j, k, t*_*k*_, *l, t*_*l*_, *m, t*_*m*_, *a, t}* at time *t*, the time evolution of the system is governed by the following forward equation:

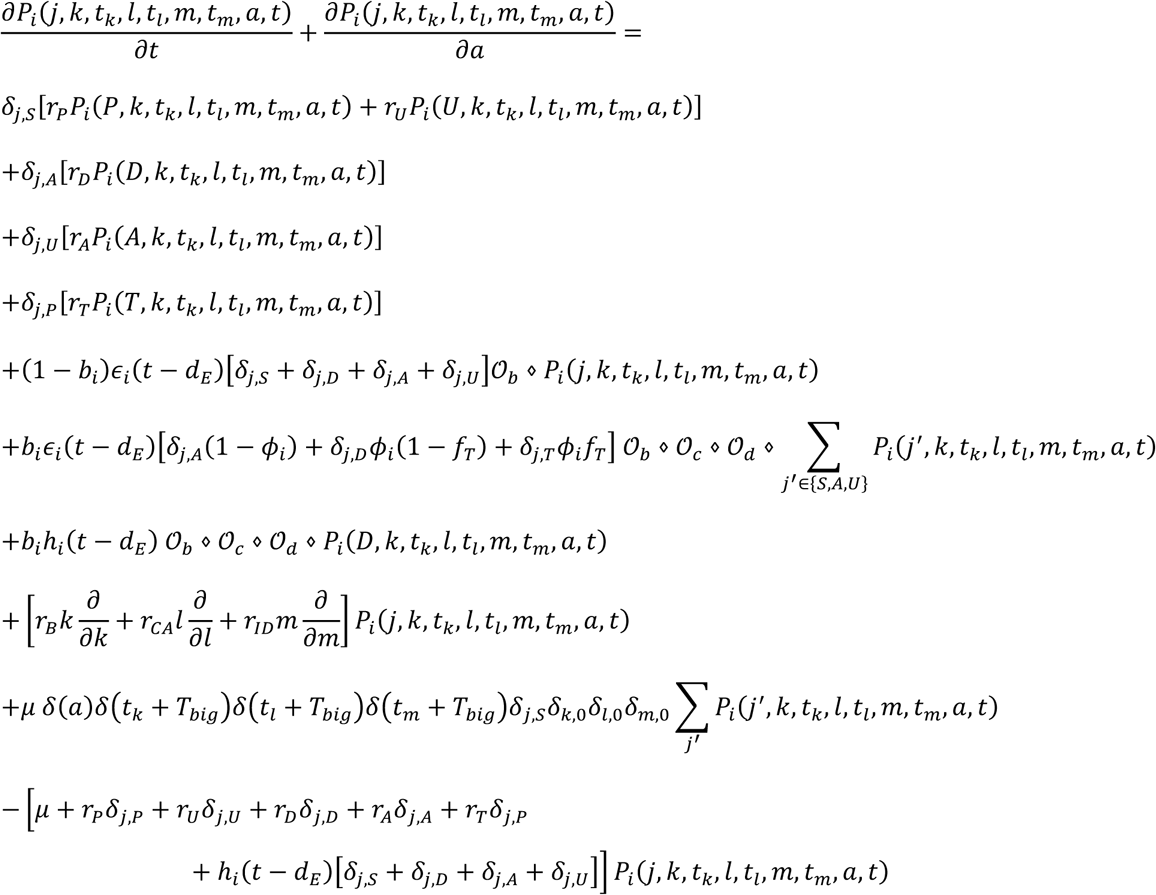

Here 𝒪_*b*_, 𝒪_*c*_ and 𝒪_*d*_ are commutative integral operators with the following action on a density *(j, k, t*_*k*_, *l, t*_*l*_, *m, t*_*m*_, *a, t)* :

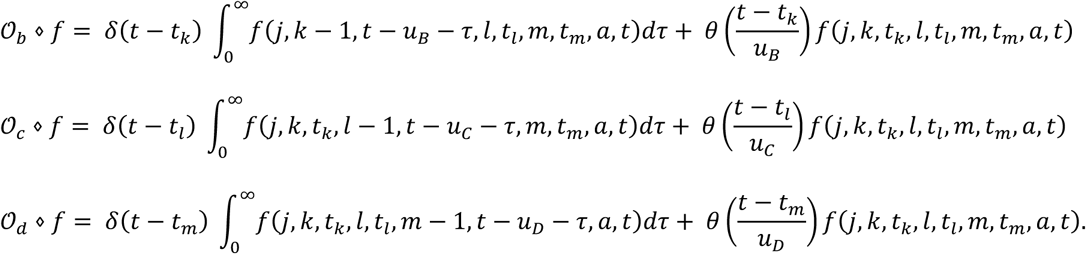

Finally, *θ(x)* is an indicator function such that *θ(x) = 1* if *x* < *1* and 0 otherwise.

For simulation, a discrete time approximation of this stochastic model was used, with a time-step of 1 day. For each individual *k, l* and *m* are set to zero at birth, while *t*_*k*_, *t*_*l*_ and *t*_*m*_ are set to a large negative value −*T*_*big*_ (to represent never having been exposed or infected, i.e. their immunity will always be boosted upon their first exposure or infection event). Each immunity term increases by 1 for an individual whenever that individual receives an infectious bite (*k*), or is infected (*l* and *m)*, if the previous boost to *k, l* and *m* occurred more than *u*_*B*_, *u*_*C*_ and *u*_*D*_ days earlier, respectively. Immunity levels decay exponentially at rate *r*_*B*_, *r*_*CA*_ and *r*_*ID*_, where *r*_*B*_, *r*_*CA*_ and *r*_*ID*_ are equal to 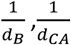 and 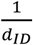 respectively.

#### Mosquito Population Dynamics

The adult stage of mosquito development was modelled individually and is similarly described in its deterministic framework before exploring its stochastic implementation. Adult mosquitoes will begin life susceptible to infection (*S*_*M*_), and will seek a blood meal on the same day they are born and every 3 days after that until the mosquito dies. Each feeding day, mosquito *i* will be exposed to a force of infection, *Λ*_*Mi*_, depending on the infection status and immunity of the human the mosquito is feeding on. The overall force of infection towards the mosquito population on a given day, *Λ*_*M*_, is thus represented by the sum of the onward infection contributions from each infected human, delayed by *d*_*g*_, delay due gametocytogenesis, which is given by

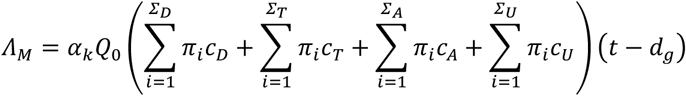

where *α*_*k*_ is the daily rate at which a mosquito takes a blood meal, *Q*_0_ is the proportion of bites that are on humans (anthropophagy) and *d*_*g*_ represents the delay from emergence of asexual blood-stage parasites to sexual gametocytes that contribute towards onward infectivity. Infected mosquitoes then pass through a latent infection stage (*E*_*M*_) that will last 10 days representing the extrinsic incubation period for the parasite (*d*_*EM*_), before becoming infectious to humans (*I*_*M*_). Infectious mosquitoes remain infectious until they die. Whenever a mosquito dies, it is replaced with a new susceptible adult mosquito. Analogously to the human model, when a new adult mosquito emerges, the day on which it dies is drawn from an exponential distribution with a transition rate of *µ*_*M*_= 0.132 days. The differential equations summarising the adult stage of mosquitoes are given by

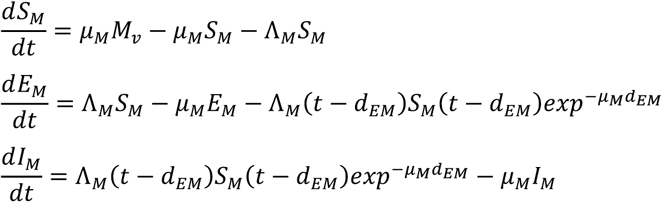

where *µ*_*M*_ is the daily death rate of adult mosquitoes, and *M*_*v*_ is the total mosquito population, i.e. *S*_*M*_ *+ E*_*M*_ *+ I*_*M*_.

#### Mosquito Stochastic Model Equations

As with the human transmission model, the full stochastic individual-based mosquito component of the model can be formally described by its Kolmogorov forward equations. As before, let *i* denote each mosquito in the population, and *j* denote their infection status. Let *δ*_*p,q*_ denote the Kronecker delta function such that it equals 1 if *p = q* and 0 otherwise. Defining *P*_*i*_(*j, t*) as the probability density function for mosquito *i* being in state {*j, t*} at time *t*, the time evolution of the system is governed by the following forward equation:

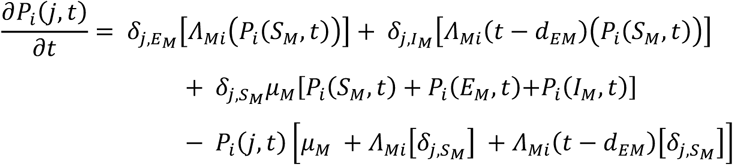

#### Seasonality and Intervention Strategies

In simulations in which no seasonality is assumed, *M*_*v*_ remains constant throughout, i.e. whenever a mosquito dies it is always replaced. When seasonality is incorporated, the maximum value that *M*_*v*_ can be oscillates with a period of 365 days. This corresponds to a change in the birth rate of mosquitoes that reflects an assumed impact upon the seasonal carrying capacity of the environment as a result of rainfall patterns upon mosquito larval stage development. In these simulations, when a mosquito dies, it will only be replaced if the current total number of mosquitoes is less than the maximum value that *M*_*v*_ can be. In simulations designed to replicate regional settings, a rainfall curve, *R(t)*, was estimated from rainfall data from 2002 to 2009 for the related first-administrative unit using the first three frequencies of the Fourier-transformed data.^6^ The seasonal total mosquito population size, *M*_*v*_*(t)*, is thus given by

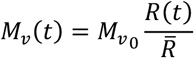

Where 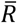 is the mean annual rainfall, and 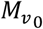 represents the seasonal harmonic mean population size.

The computational constraints introduced by modelling individual mosquitoes and parasite population genetic dynamics necessitated modelling intervention strategies indirectly. This was handled by assuming that an introduction of intervention leads to a decrease in the average age of the mosquito population throughout the duration of the intervention due to an increased mortality rate. As a result, the average age reflects a new composite mortality rate due to both interventions and external causes. Similarly it leads to an increase in *Q*_0_ to reflect mosquitoes that are repelled as a result of interventions but do not die. The daily rate of change to these parameters in response to ITN and IRS coverage is calculated using an equivalent deterministic version of the earlier model that included interventions,^1^ before being introduced as a time-dependent variable within the stochastic model.

### Parasite Dynamics

#### Parasite Genetic Barcode

Parasites are modelled as discrete populations as a result of an infection event associated with a mosquito or a human. Each asexual parasite is characterised by one genetic barcode, which contains information relating to 24-SNPs distributed across the parasite genome. These SNPs represent an increasingly used general SNP-based molecular barcode that has been used for the identification and tracking of *P. falciparum* clones.^7^ Sexual stages of the parasite lifecycle within the mosquito are represented by both a female and male barcode, thus defining the range of recombinants that could be produced. The within human parasite dynamics and model considerations are discussed first before exploring the within mosquito parasite life cycle and associated modelling implications. A schematic overview of the modelled parasite lifecycle stages is shown in Diagram 2.

In simulations modelling identity-by-descent (IBD), we extend the barcode to consider 24 “identity-loci”. An identity loci can take any integer value required, allowing true identities to be compared. In the SNP-loci barcode, each loci can only be 0 or 1, representing the minor and major allele for that barcode loci.

#### Within Human Parasite Dynamics

During a successful mosquito to human infection event, a number of asexual parasite barcodes are introduced into the human, which may be observed in the ensuing gametocyte genotypes when considering onward infectiousness from humans to mosquitoes. If the individual’s pre-erythrocytic immunity was boosted in the last *u*_*B*_ days no new parasite barcodes will be passed to the individual, otherwise more than one different asexual parasite barcode that will be observed in the ensuing gametocyte genotypes may be introduced during an infection event, representing cotransmission of genetically related parasites (if the mosquito was infected with more than one sporozoite genotype). The precise distribution describing the number of genotypes is unknown,^8^ but the mean number of sporozoites within an inoculation event is well characterised by a geometric distribution with mean equal to 10. The geometric mean will then be used to estimate the proportion of sporozoites that are successful, *ξ*, which yields the maximum number of successful sporozoites in an individual with no pre-erythrocytic immunity. If this number is less than 1, then a new total number of sporozoites is drawn until the maximum number of sporozoites after incorporating *ξ* is greater than 0. The observed number of successful sporozoites is then calculated by conducting Bernoulli trials for all but one of the successful sporozoites (as we assume one has to survive to found the infection) to see if they are successful, calculated using the individual’s probability of infection, *b*_*i*_. In summary this can be written as:

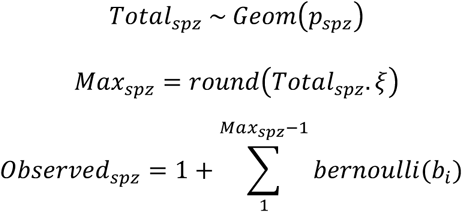

There is no assumed maximum number of parasites, with individuals assumed to clear strains on the day that they would have moved from a subpatent infection to susceptible for the strain considered, i.e. each acquired strain follows an assumed trajectory in parasitaemia representative of a normal infection cycle, i.e. with a mean duration of infectiousness equal to *d*_*A*_ + *d*_*U*_. Acquired strains can thus move “infection state” independently of the human’s infection state. For example, a given individual is infected on day 0 and develops an asymptomatic infection. The individual is scheduled to become subpatent on day 200, but they were bitten on day 150 and developed clinical symptoms and moved to state D. When this happens, the parasite density of the strain acquired on day 0 does not change and this strain will become a subpatent strain on day 200. After day 200, its probability of being onwardly transmitted is thus equal to *c*_*U*_. After the parasite has moved to become a subpatent strain, the day at which the strain would have been cleared, i.e. the individual would have moved from state U to S if they had not been superinfected, is drawn and assigned to the parasite. On this drawn day the subpatent parasite strain is assumed to have been cleared. By tracking parasites in this way we are able to track the relative parasitemias of each acquired strain, enabling more accurate sampling of within host parasite genetic diversity when passing on gametocytes to mosquitoes as well as enabling an equilibrium between clearing old strains and acquiring new strains, which represents the multiplicity of infection. This is shown in the schematic below (Diagram 2), which also details the key features of the barcode.

#### Within mosquito parasite dynamics

When a mosquito is infected, we sample from a zero-truncated negative binomial distribution that describes the distribution of oocysts that form from a feeding event. The choice of a zero truncated negative binomial represents the increasingly identified zero-inflated negative binomial that describes the relationship between oocyst prevalence and mean oocysts per mosquito in SMFA studies.^9–11^ The related negative binomial distribution for the distribution of oocysts is given by

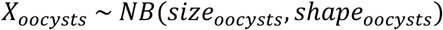

where *X*_*oocysts*_ represents the number of oocysts that will be formed, with mean equal to 2.5 and a shape equal to 1, which captures the mean and range of oocysts observed in natural *P. falciparum* infections^9,10,12^ For each oocyst formed, two barcodes are sampled from the infected host representing the female and male gametes that led to the oocysts formation. These two barcodes will result in up to 4 different potential genotypes (reflecting the immediate two step meiotic division that takes place after zygote formation) represented within the sporozoite population within the oocyst. When an infectious mosquito seeks a blood meal and leads to an onward infection, a value for *Observed*_*spz*_ is sampled. The oocyst source for each onward infection within a coinfection is sampled from oocysts that have ruptured, i.e. the infection event that led to the oocyst occurred more than 10 days earlier. At this point recombination is simulated by randomly choosing either the male or female allele at each SNP position in the barcode. The random sampling in this represents the assumed independent segregation events resulting from the absence of genetic linkage between barcode SNP positions. Once a recombinant has been simulated it is stored and associated with the oocyst from which it came. If the same oocyst is chosen to lead to an additional infection, then the previously generated recombinant has a 25% chance of being onwardly transmitted and there is a 75% chance that a new recombinant is generated and subsequently saved. This process will continue in ensuing onward infection events that result from this oocyst until four recombinants have been simulated, at which point they each have a 25% chance of being onwardly transmitted. The above thus introduces an assumption that sporozoites will remain onwardly-transmissible for the remainder of the mosquito’s life, with no effect upon their relative probability of being onwardly transmitted in relation to sporozoites that resulted from a more recently ruptured oocyst.

**Diagram 2:**
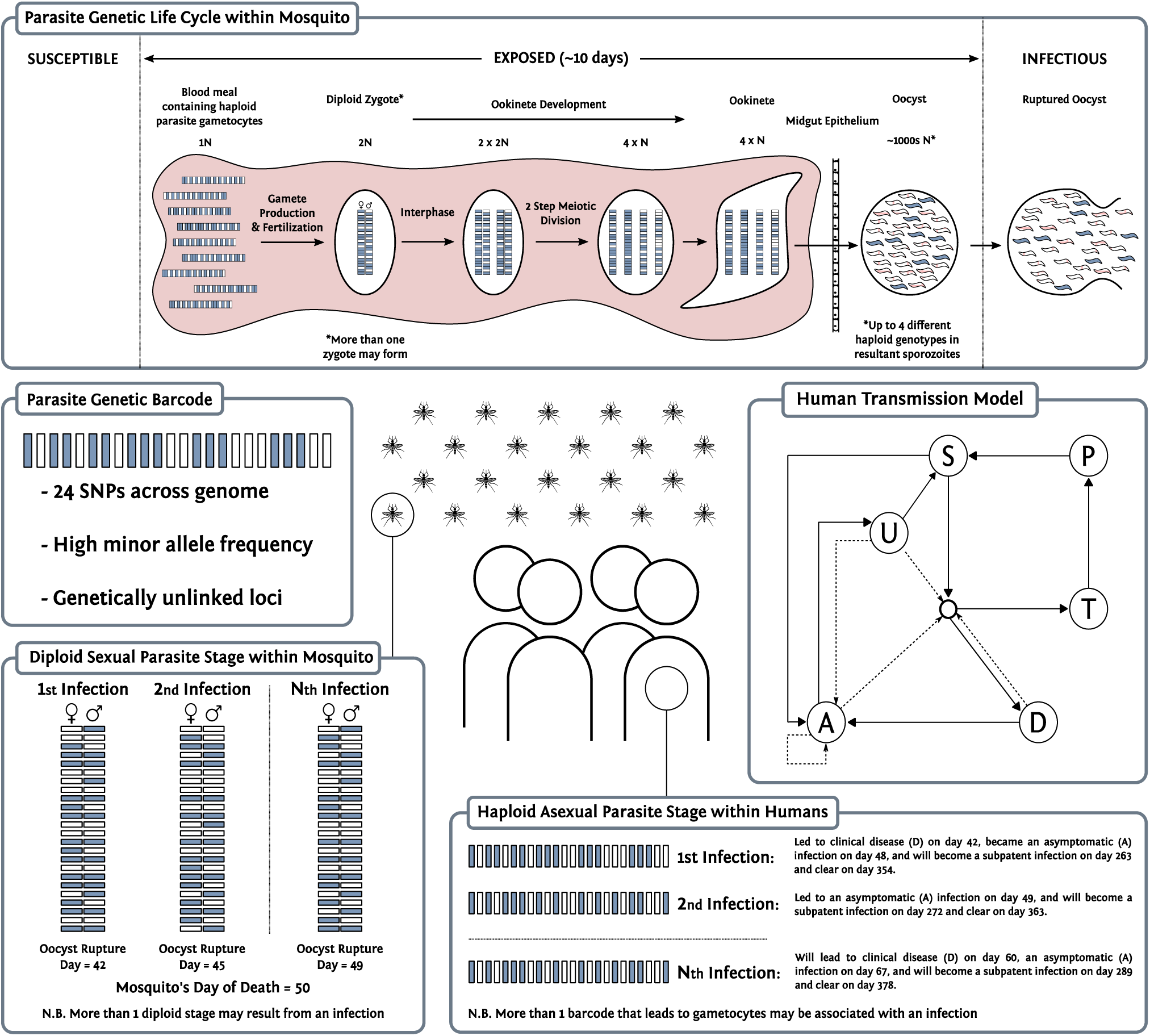
Parasite Dynamics within the transmission model. Individual mosquitoes are tracked, which allows for recombination to be modelled explicitly. Populations of parasite clones are tracked, and multiple oocysts are able to be formed from a feeding event, as well as multiple genetically distinct sporozoites onwardly transmitted. A “barcode” is associated with each parasite clone and can either represent biallelic SNPs, or unique identities that allow IBD to be calculated.

#### Importation Rate

The non-spatial, closed population nature of the model will result in the eventual fixation of a single genetic barcode. As such, when conducting simulations designed to replicate regional settings, an estimate of the importation rate was calculated, yielding to a daily probability that an infection is due to an imported case. The importation rate represents the sum of two different flows of infection into a regional setting:

A. Individuals who are infected outside the region while travelling to and from other areas
B. Visiting travellers from outside the region who infect mosquitoes within the admin unit

These two process are incorporated at the same stage within the model, whereby there is a temporally dependent daily probability that a generated recombinant genotype is due to an importation as follows

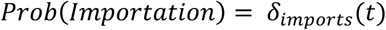

where *δ*_*imports*_*(t)* is the population proportion of new infections resulting from importations on a given day. This parameter changes over time to reflect changes in regional seasonality (both within the region and neighbouring regions), and different rates of change in malaria prevalence across neighbouring regions.^13^ If the recombinant is due to an importation, then a random barcode is produced and passed on. This barcode will also be stored and associated with an oocyst within the mosquito considered if it was probabilistically determined to be due to the second flow of importation defined above (B), determined by the ratio of these two flows of infection. Predicted rates of the two flows of infection above are calculated for each year between 2000 and 2015 using a fitted gravity model of human mobility.^14^

### Model Parameter Values

**Table 1:**
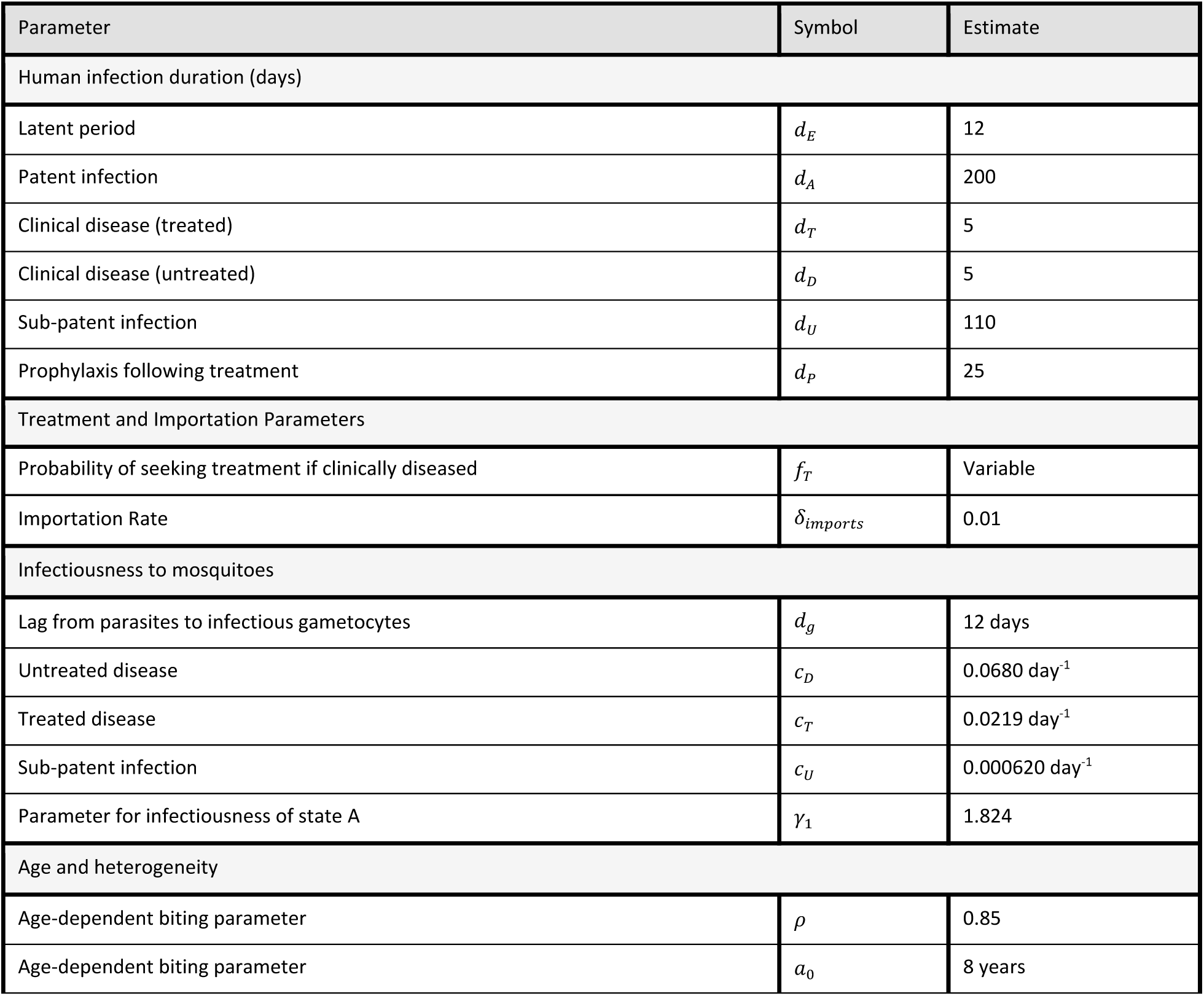
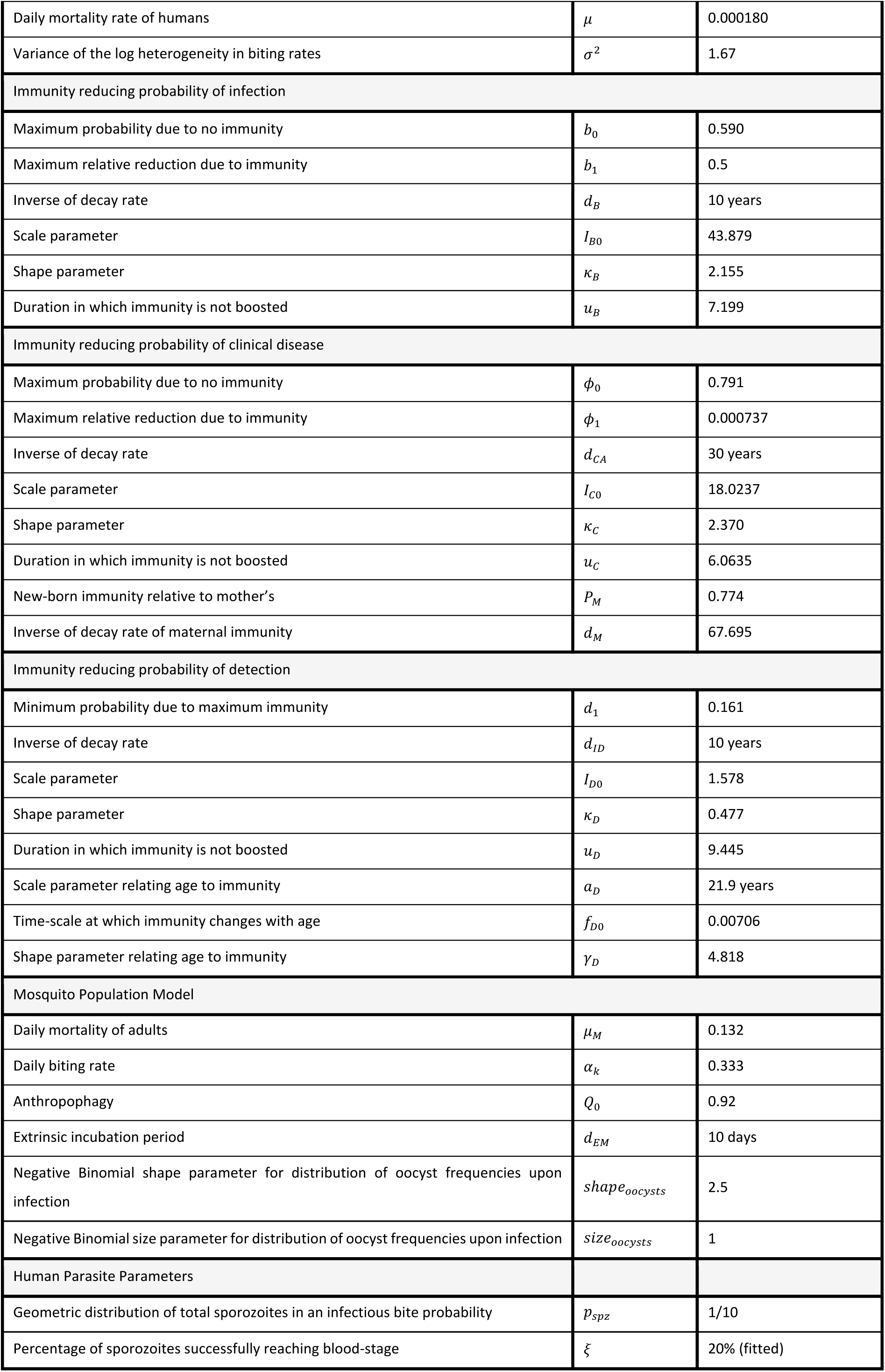
Parameter estimates used within the model were taken from Griffin et al. 2014,^3^ 2015^2^ and 2016^1^.

| Parameter | Symbol | Estimate |
| --- | --- | --- |
| Human infection duration (days) |  |  |
| Latent period | $d_E$ | 12 |
| Patent infection | $d_A$ | 200 |
| Clinical disease (treated) | $d_T$ | 5 |
| Clinical disease (untreated) | $d_D$ | 5 |
| Sub-patent infection | $d_U$ | 110 |
| Prophylaxis following treatment | $d_P$ | 25 |
| Treatment and Importation Parameters |  |  |
| Probability of seeking treatment if clinically diseased | $f_T$ | Variable |
| Importation Rate | $\delta_{imports}$ | 0.01 |
| Infectiousness to mosquitoes |  |  |
| Lag from parasites to infectious gametocytes | $d_g$ | 12 days |
| Untreated disease | $c_D$ | 0.0680 day<sup>-1</sup> |
| Treated disease | $c_T$ | 0.0219 day<sup>-1</sup> |
| Sub-patent infection | $c_U$ | 0.000620 day<sup>-1</sup> |
| Parameter for infectiousness of state A | $\gamma_1$ | 1.824 |
| Age and heterogeneity |  |  |
| Age-dependent biting parameter | $\rho$ | 0.85 |
| Age-dependent biting parameter | $a_0$ | 8 years |
| Daily mortality rate of humans | $\mu$ | 0.000180 |
| Variance of the log heterogeneity in biting rates | $\sigma^2$ | 1.67 |
| Immunity reducing probability of infection |  |  |
| Maximum probability due to no immunity | $b_0$ | 0.590 |
| Maximum relative reduction due to immunity | $b_1$ | 0.5 |
| Inverse of decay rate | $d_B$ | 10 years |
| Scale parameter | $I_{B0}$ | 43.879 |
| Shape parameter | $\kappa_B$ | 2.155 |
| Duration in which immunity is not boosted | $u_B$ | 7.199 |
| Immunity reducing probability of clinical disease |  |  |
| Maximum probability due to no immunity | $\phi_0$ | 0.791 |
| Maximum relative reduction due to immunity | $\phi_1$ | 0.000737 |
| Inverse of decay rate | $d_{CA}$ | 30 years |
| Scale parameter | $I_{C0}$ | 18.0237 |
| Shape parameter | $\kappa_C$ | 2.370 |
| Duration in which immunity is not boosted | $u_C$ | 6.0635 |
| New-born immunity relative to mother's | $P_M$ | 0.774 |
| Inverse of decay rate of maternal immunity | $d_M$ | 67.695 |
| Immunity reducing probability of detection |  |  |
| Minimum probability due to maximum immunity | $d_1$ | 0.161 |
| Inverse of decay rate | $d_{ID}$ | 10 years |
| Scale parameter | $I_{D0}$ | 1.578 |
| Shape parameter | $\kappa_D$ | 0.477 |
| Duration in which immunity is not boosted | $u_D$ | 9.445 |
| Scale parameter relating age to immunity | $a_D$ | 21.9 years |
| Time-scale at which immunity changes with age | $f_{D0}$ | 0.00706 |
| Shape parameter relating age to immunity | $\gamma_D$ | 4.818 |
| Mosquito Population Model |  |  |
| Daily mortality of adults | $\mu_M$ | 0.132 |
| Daily biting rate | $\alpha_k$ | 0.333 |
| Anthropophagy | $Q_0$ | 0.92 |
| Extrinsic incubation period | $d_{EM}$ | 10 days |
| Negative Binomial shape parameter for distribution of oocyst frequencies upon infection | $shape_{oocysts}$ | 2.5 |
| Negative Binomial size parameter for distribution of oocyst frequencies upon infection | $size_{oocysts}$ | 1 |
| Human Parasite Parameters |  |  |
| Geometric distribution of total sporozoites in an infectious bite probability | $p_{spz}$ | 1/10 |
| Percentage of sporozoites successfully reaching blood-stage | $\xi$ | 20% (fitted) |

